# miRNA Sensor HuR Compartmentalizes Ago2-Uncoupled miRNAs to Lipid Droplets to Buffer miRNA Activity in Mammalian Cells

**DOI:** 10.64898/2026.01.10.698809

**Authors:** Sreemoyee Chakraborty, Diptankar Bandyopadhyay, Kamalika Mukherjee, Suvendra N. Bhattacharyya

## Abstract

miRNA activity must be optimized to ensure target gene expression in mammalian cells in response to specific needs. How and when miRNA activity is regulated remains an open question in gene expression regulation, with limited information on how cellular machinery senses miRNA levels to regulate miRNA expression and activity. Because miRNAs are stable molecules that can be reversibly used, the existence of a miRNA-sensing mechanism in mammalian cells has been anticipated. With ectopic expression of miRNAs in mammalian cells, we found a dose-dependent miRNA buffering mechanism in which miRNA export and its storage on lipid droplets are essential for miRNA activity optimization in mammalian cells. We found that the miRNA-binding protein HuR, known for its role in miRNA export, has a dual function in mammalian cells. HuR uncouples miRNAs from Ago2 to facilitate association of lipid droplets with nonfunctional miRNAs, a process that gets augmented in cells with high lipid droplet content and restricted extracellular export. Thus, HuR acts as a miRNA sensor, and the optimal activity and abundance of HuR regulate miRNA storage on lipid droplets or its export to maintain cellular miRNA homeostasis, thereby preventing detrimental effects of an excess miRNA pool in mammalian cells. Thus, targeting miRNA export or lipid droplet association is an important strategy for buffering cellular miRNA levels.

**Importance:** To ensure that target gene expression in mammalian cells is appropriately regulated, miRNA activity needs to be carefully fine-tuned. Although miRNAs are stable and can be reused, the precise mechanisms by which their levels are sensed and regulated remain unclear. Our research uncovered a fascinating dose-dependent miRNA buffering process that involves export and storage of miRNAs on lipid droplets—essential steps for precise regulation. We also found that HuR, a protein that facilitates miRNA export, plays a key role by uncoupling miRNAs from Ago2 and promoting their association with lipid droplets in cells with abundant lipid droplets and limited extracellular export capacity. Overall, HuR acts as a miRNA sensor, helping to balance miRNA storage and export to maintain cellular homeostasis and prevent the harmful effects of miRNA overactivity.

## Introduction

miRNAs are small regulatory RNAs of 22 nt that repress expression of the encoded proteins by binding to target mRNAs(1, 2). The activity of miRNAs must be tightly regulated, as their abundance and activity play a significant role in determining cellular fate upon exposure to external cues(3). Most miRNA-regulated processes require an active miRNA complexed with Ago proteins to bind to and induce the degradation of target mRNAs, as miRNA-mediated regulation occurs primarily at the post-transcriptional level(3, 4).

Previous studies have highlighted the pivotal roles of subcellular compartments in regulating miRNA levels and activity in mammalian cells(5). Polysomes attached to the endoplasmic reticulum have been identified as sites of miRNA-target mRNA interaction(6, 7), a process controlled by mitochondrial activity and mTORC1(8, 9). We have also previously identified how repressed mRNAs are targeted to endosomes(10), where miRNAs are decoupled from target mRNAs and either recycled via lysosomal pathways(11) or acted on by miRNA-binding proteins such as HuR to facilitate Ago2-miRNA unbinding. HuR-controlled miRNA export(12), a process also supported by HuR-interacting membrane proteins STX5(13) and RalA GTPase(14). The RNA-processing body targeting of miRNAs and repressed messages constitutes a counteracting mechanism that prevents HuR-mediated uncoupling and export of miRNAs(15). In contrast, the GW182 protein, an RNA-processing body component, ensures P-body targeting of miRNAs, thereby facilitating P-body-mediated storage or degradation of miRNA target messages(15). The regulation of miRNA activity is essential for optimal expression of target genes, and derepression of miRNA-repressed messages can contribute to stress responses(16), immune activation(17), regulation of cancer cell proliferation(15), and neuronal differentiation(18, 19).

It is an essential question in gene expression regulation research: how miRNA activity is regulated. The transcriptional regulation of miRNAs by specific transcription factors and the processing of pre- and pri-miRNAs in the nucleus and cytoplasm are essential steps in miRNA biogenesis(20, 21). However, given the stability of miRNAs relative to other RNAs and the reported exchange of miRNAs between mammalian cells via extracellular vesicles, several mechanisms can regulate miRNA function at the post-transcriptional level. Extracellular export is considered a key mechanism of miRNA regulation in mammalian cells (22, 23), thereby restricting miRNA levels on demand. However, it appears to be an irreversible mechanism for controlling miRNA expression, as are the reported miRNA degradation machineries, which may also determine miRNA levels in mammalian cells(23, 24). However, given the reversible nature of miRNA action on its target(16), rapid and reversible buffering of miRNA activity is anticipated in cells primed for the genesis of excess miRNAs, as increased miRNA biogenesis machinery activity during stress-response development can drive excess miRNA production(8, 25, 26). On the contrary, the reversible regulation of miRNA activity is equally important for controlling events during neuronal differentiation and for local translational processes reported in brain cells(27, 28, 29).

Excess miRNAs that require buffering should be sensed by specific RNA-binding proteins, which regulate miRNA activity without detrimental effects on cellular homeostasis, thereby responding to external cues. Interestingly, in hepatic cells exposed to high lipid levels, miRNA activity must be rapidly regulated to counteract lipotoxicity, and cells export Dicer1, the key protein required for miRNA biogenesis, to prevent excess production of miR-122, a miRNA implicated in cell death upon exposure to high lipids(30).

Following cells with high lipid levels and increased lipid droplet (LD) numbers(30), we found an association between miRNAs and lipid droplets in hepatic cells. Lipid droplets are known for their lipid storage function, and recent evidence suggests they may have a broader role, as they are involved in processes beyond lipid metabolism and energy homeostasis in mammalian cells(31, 32). We hypothesize that LDs play a pivotal role in miRNA storage, which is required for miRNA buffering in mammalian cells. Increased expression of mature miRNAs is balanced by EV-mediated export and miRNA storage in LDs, thereby preventing excessive accumulation of functional miRNAs. We found that HuR plays a dual role in this buffering process. HuR, known to facilitate miRNA export via extracellular vesicles, also buffers miRNA levels by promoting miRNA storage in LDs. HuR does so by uncoupling miRNAs from Ago proteins. HuR acts as a miRNA sensor that, in response to cellular demand, modulates miRNA storage and export to reduce miRNA activity and prevent cellular damage under specific cellular needs, such as avoiding lipotoxicity-related death.

## Results

### miRNA buffering ensures the homeostasis of miRNA levels in human cells

Two main pathways that regulate miRNA levels in mammalian cells are miRNA biogenesis by Dicer1/TRBP(33) and miRNA export via extracellular vesicles (EVs) into the extracellular space(34). There is also evidence of post-transcriptional modifications that enable miRNAs to be selectively degraded by specific exonucleases, thereby controlling miRNA levels in mammalian cells(35). How mammalian cells sense miRNA levels and how their intracellular concentrations are regulated to balance the functional miRNA pool remain largely unanswered(36). To investigate this, we examined the effects of introducing high levels of specific miRNAs into cells and tracked their fate as their cellular concentration increased. We sought to increase cellular miRNA levels stepwise to document their effects on miRNA levels and activity without altering pre- and pri-miRNA levels. We transfected HeLa cells with increasing amounts of a miR-122 mimic, a double-stranded form of this liver-specific miRNA that is otherwise not expressed in HeLa cells. Functional miR-122 production from the mimic is Drosha/Dicer-independent(37, 38). We monitored mature miR-122 levels in the cytoplasm and extracellular space, with and without GW4869, to assess the effect of blocking miRNA export on intracellular miRNA content. GW4869, a neutral sphingomyelinase inhibitor, is known to block extracellular vesicle (EV) or exosome-mediated miRNA export (**Fig. 1A**)(39). As expected, we observed increased export of miR-122 as miRNA content of EVs isolated from HeLa cells transfected with the mimic, rising with higher mimic concentrations used for transfection (**Fig. 1B**). Cellular miR-122 levels gradually increased with higher mimic concentrations, but decreased at 200 nM, an effect prevented by GW4869, which is known to inhibit miRNA export. This indicates that the miRNA buffering capacity of a human cell is concentration-dependent, reaching a threshold at 100 nM for miR-122, which then decreases at 200 nM, the mimic concentration used for transfection. This suggests that export via EVs may play a key role in miRNA homeostasis and loss in the presence of GW4869 (**Fig. 1C**). Counting on the amount of miR-122 used for transfection and the Ct values we observed, it is approximately 20,000 and 23,000 copy of miR-122 per cells when 50 and 100 nM miR-122 mimic were used for transfection which drops to 16,000 miR-122 per cells at 200 nM mimic concentration in control cells without GW4869 treatment.

**Figure 1.**
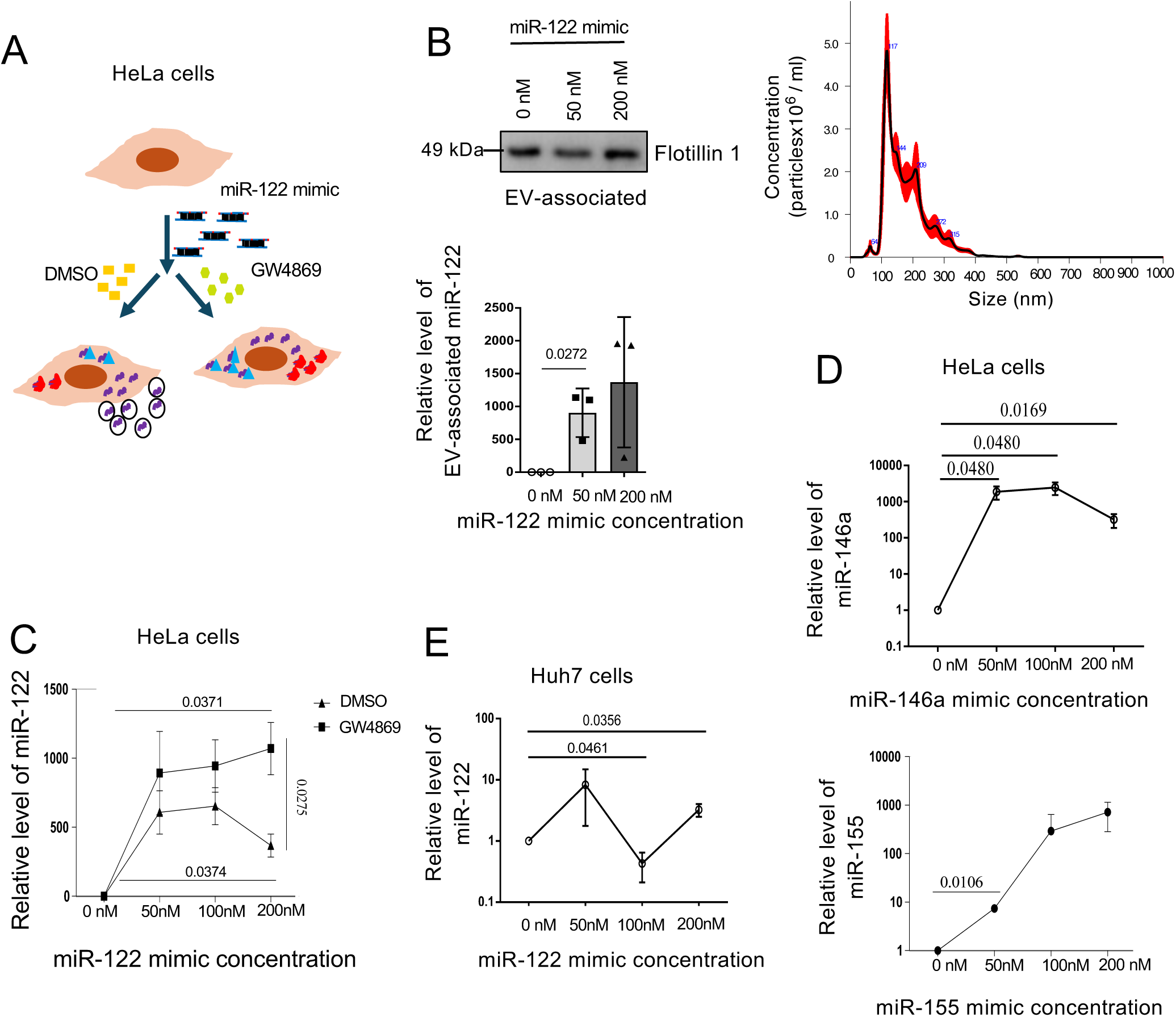
miRNA buffering in mammalian cells is controlled by extracellular export. A. Schematic of the experimental methodology used to study miRNA buffering in HeLa cells. B. Quantification of miR-122 in extracellular vesicles isolated from HeLa cells transfected with increasing concentrations of miR-122 mimic (n=3). Data were normalized to the EV marker Flotillin1 protein levels. The presence of Flotillin1 in the Western blot of the EV samples confirmed effective EV isolation. The right-hand panel depicts the NTA data obtained with EVs isolated from HeLa cells. C. Graph depicting the cellular miR-122 levels quantified by qRT-PCR and value normalized against U6 RNA in DMSO control and GW4869 (10μM, 24h) treated HeLa cells transfected with 200nM miR-122 mimic (n=3). D. Relative levels of cellular miR-146a and miR-155 in HeLa cells transfected with increasing concentrations of the respective miRNA mimics after 48 h of transfection. qRT-PCR was used for estimation of miRNAs normalized against U6 RNA levels (n=3). E. Effect of miR-122 mimic transfection on cellular miRNA levels measured by qRT-PCR in Huh7 cells. U6 RNA was used for normalization for experiments done in triplicate. Data are presented as SEM ± SD, and P values are reported in the respective panels tested for statistical significance. The positions of the molecular weight markers are shown in the Western blot panel. P-values were calculated by a two-tailed paired t-test in most of the experiments unless mentioned otherwise. The fold change of miRNA was calculated by 2^−ΔCt^ method.

Thus, the buffering effect is impaired if miRNA export is blocked. The miRNA buffering event is not limited to miR-122; similar patterns were observed with miR-146a and miR-155 in HeLa cells, in which cellular levels declined after reaching a threshold. Interestingly, these thresholds may differ for different miRNAs (**Fig. 1D**). In hepatic cells, where a pre-existing endogenous pool of miR-122 is already present [16,000 copies/cell(12)], the cellular thresholds for miR-122 were found to be biphasic, with a reduction in miR-122 levels even at lower concentrations (100 nM), which partially recovered at 200 nM (**Fig. 1E**), suggesting a dual mode of miRNA buffering may be operative in hepatic cells for miR-122. We speculate that miRNA buffering may be mediated by lysosomal pathways that degrade excess miRNAs when required. The contribution of the lysosomal pathway to buffering miRNAs has been ruled out by experiments conducted in the presence of Bafilomycin, which selectively blocks lysosomal targeting of endosomal content and its degradation, manifested by accumulation of LC3B II, but without any effect on miRNA content at 200nM miR-122 mimic transfection condition, suggesting that the lysosomal pathway may have an insignificant impact on miRNA buffering observed (**Fig. S1**).

### Cellular miRNAs accumulated beyond the cellular threshold remain non-functional in mammalian cells

Do miRNAs that accumulate after the cellular threshold is reached remain functional in mammalian cells, or do they stay dormant, localized in specific subcellular compartments or phase-separated structures, as previously observed in amyloid-exposed glial cells(11)? To address this question, we measured miR-122 repressive activity in HeLa cells following transfection with the miR-122 mimic. We observed that the specific activity of miR-122, measured as the fold repression of target reporter mRNAs per unit miRNA, decreased with increasing miR-122 mimic concentration used for transfection (**Fig. 2A and 2B**). Blocking the extracellular export of miR-122 with GW4869, rather than increasing miRNA activity, reduced miRNA-mediated reporter mRNA repression, consistent with the idea that accumulated miRNAs are nonfunctional once they reach the threshold (**Fig. 2C**). A similar observation was also noted for the endogenous target of miR-122, as the CAT1 and Aldolase mRNA levels do not show further repression at 200 nM in the presence of GW4869 (**Fig. 2D**). The apparent decrease in cellular miRNA activity caused by a failure of the miRNA machinery in the presence of GW4869 and at miRNA concentrations above the cellular threshold could explain this observation. Supporting this hypothesis, levels of endogenous let-7a and miR-21 change with miR-122 transfection, and the repressive effect of let-7a on the Renilla reporter containing let-7a binding sites shows a similar pattern in HeLa cells treated with 200 nM miR-122 and GW4869 (**Fig. 2E**). This suggests that the miRNA machinery is overwhelmed when excess miRNAs are present without an miRNA-export mechanism to remove the extra miRNAs.

**Fig. 2.**
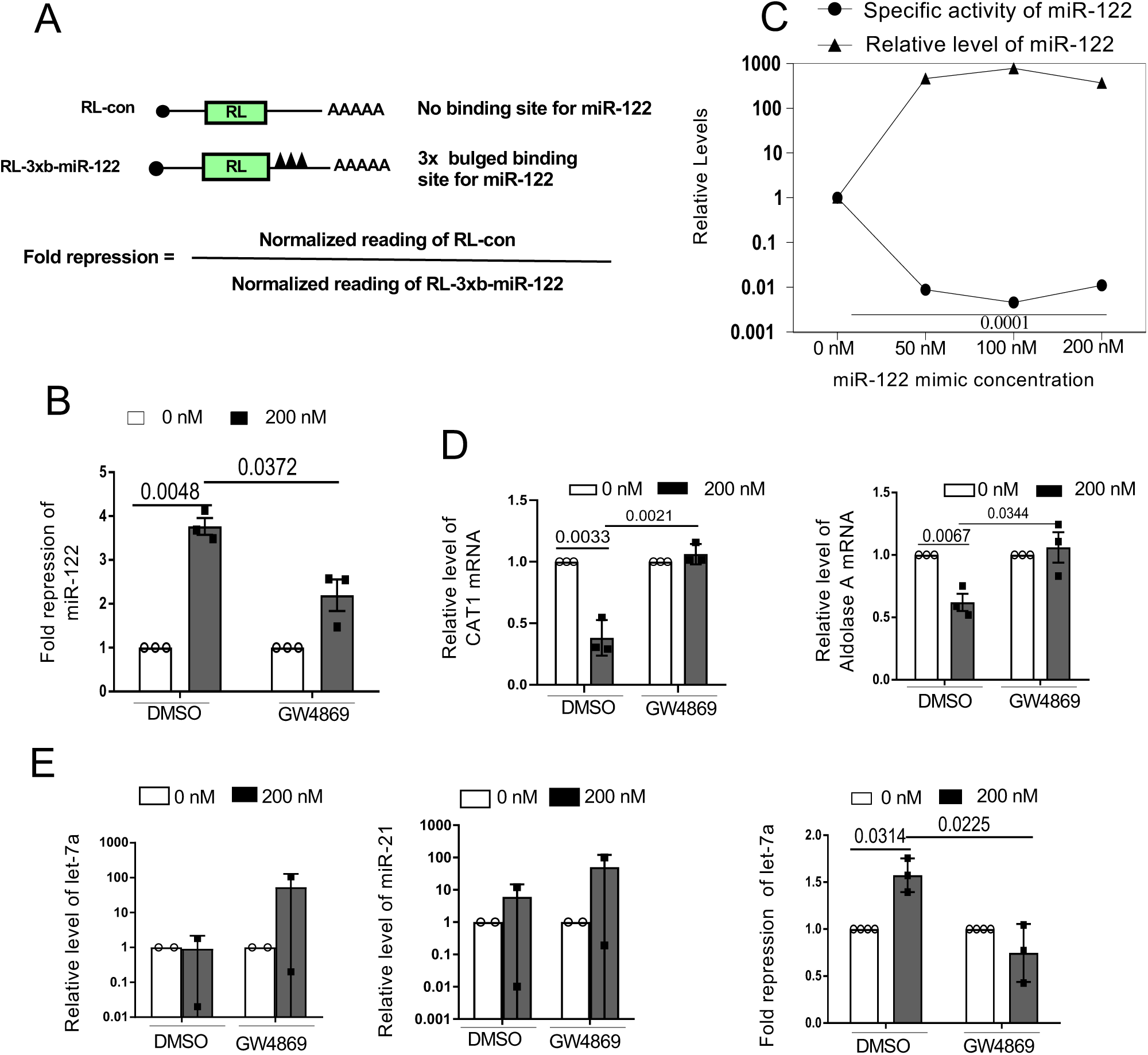
miRNAs accumulated in cells blocked for export are repression-defective. A. A schematic of the Reporter mRNAs used for measuring miRNA activity in HeLa cells. Fold repression is calculated as the ratio of firefly luciferase-normalized Renilla reporter expression for miR-122 to that of the control Renilla reporter lacking miRNA-binding sites. B. Fold repression of Renilla reporter of miR-122 in control and GW4869-treated cells, transfected with 0 or 200nM of miR-122 mimic. Experiments were done in triplicate. C. Fold repression of reporter mRNA with increasing miRNA mimic concentrations has been used to calculate the specific activity of miR-122 in HeLa cells (Specific activity = fold repression per unit miRNA). The level of cellular miRNAs is plotted as a function of concentration. Data from three experimental replicates are used in the calculations. D. Relative levels of miR-122 endogenous targets, CAT-1 and Aldolase mRNA levels, were measured in control (DMSO) and GW4869-treated HeLa cells transfected with 200nM miR-122 mimic. E. Effect of miR-122 mimic transfection (200nM) on let-7a and miR-21 miRNA levels (n=2) and let-7a repressive activity on the Renilla let-7a reporter in HeLa cells treated with DMSO or GW4869 (right panel, n=3). Data are presented as SEM ± SD, and P values are reported in the respective panels tested for statistical significance. P-values were calculated by a two-tailed paired t-test in most of the experiments unless mentioned otherwise. The fold change of miRNA was calculated by 2^−ΔCt^ method.

### Lipid droplets store miRNAs and contribute to miRNA buffering in mammalian hepatic cells

The storage of mRNA in its non-functional form has been reported before in RNA processing bodies(11). With the anticipation that P-bodies contribute to the storage of non-functional miRNAs when miRNAs are in excess, it was an exciting hypothesis to investigate. We did not observe any change in miRNA localization with Dcp1a, although we observed a non-significant increase in PB-number in cells transfected with 200 nM miR-122 mimic. The transfecting miRNAs were observed as foci proximal to P-bodies with no marked overlap detected (**Fig.S2 A and B**).

We were interested to know the location of these miRNA bodies that are observed in miR-122 mimic-transfected cells. In a lipid-abundant state, liver cells generate Lipid droplets, a single membrane organelle designated as lipid storage sites, to store additional lipids(40). High lipid levels reduce miR-122 activity, and to prevent the genesis of new miRNAs, Dicer1 is exported, thereby preventing a lipotoxic state(30). It has been reported that miR-122 prevents lipid droplet (LD) formation(41), while other studies suggest a positive correlation between miR-122 and lipid droplets(42). Using the hydrophobic lipid-binding BODIPY 493/503(43), which binds to hydrophobic lipids in LD, we detected an increase in LD number in HeLa cells transfected with 200nM miR-122. We observed colocalization of miR-122 with lipid droplets using 200nM miRNA mimics for transfection, and the number of LDs increased significantly at 200nM but not at 50 nM concentration (**Fig.3 A-C**).

**Figure 3.**
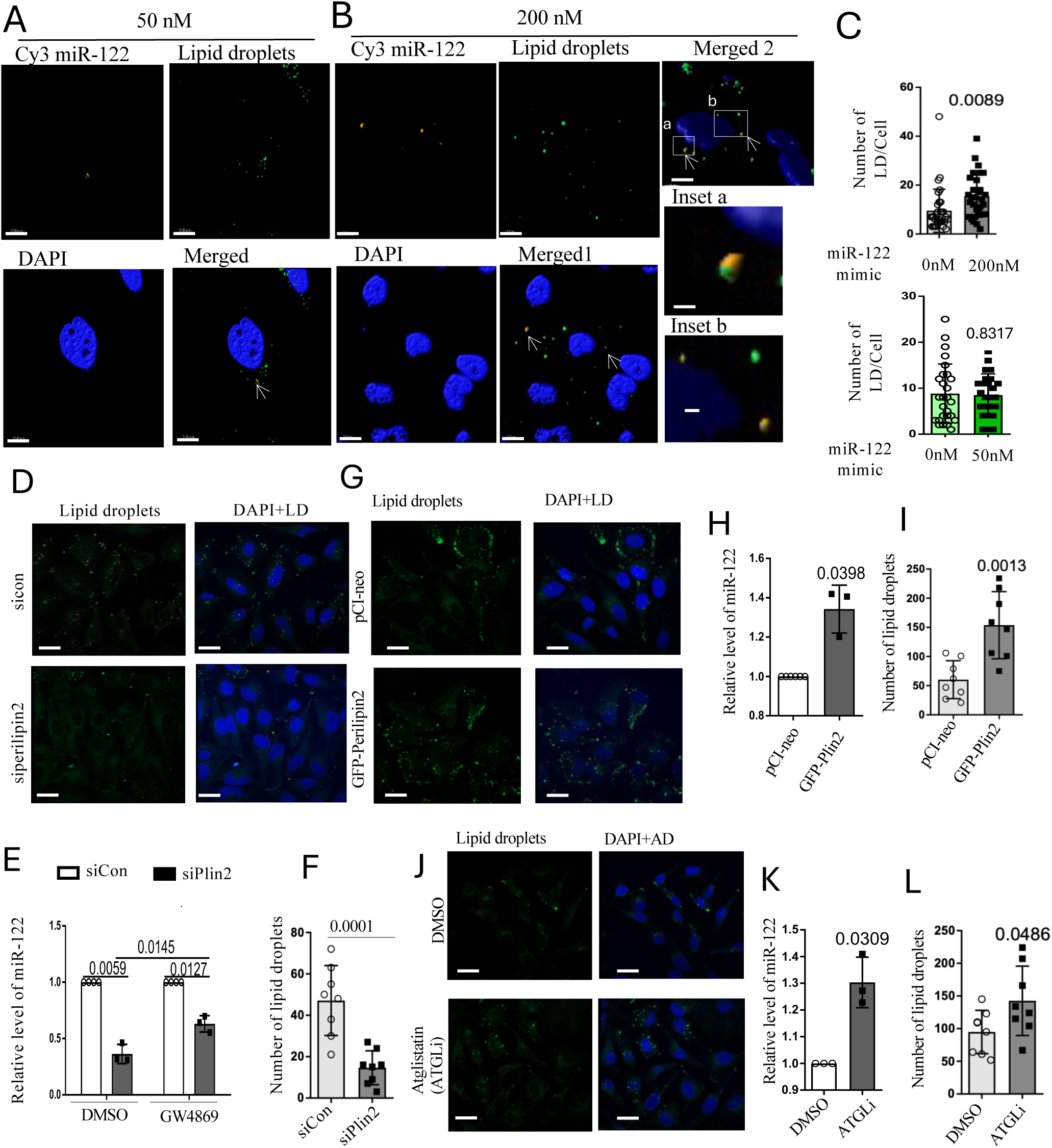
Lipid droplets control cellular miRNA content. A-B. Effect of miR-122 on the number of lipid droplets in HeLa cells. Image of the Cy-3 labeled miR-122 (red) used for transfection (50nm; A) and (200 nM; B), and lipid droplets visualized with BODIPY 493/503 dye (green). DAPI stains the nucleus. Scale bars are 10μm and 2 μm (zoomed panel). Merged 1 and 2 showing different cells in 200nM miR-122 mimic transfection conditions. In Merged 2, the a and b parts marked are zoomed to show colocalization of miRNA and LD. A. Effect of miR-122 mimic transfection ( 50 or 200 nM) on lipid droplet (LD) number. Quantification of LDs number (>100nm) observed in confocal images of the cells stained for BODIPY 493/503 dye. A minimum of 20 cells was counted for each set. D-E. Effect of Perilipin 2 depletion by siRNAs in HeLa cells on LD (D) and miRNA levels (E). The SiRNA mixture targeting Perilipin 2 was used to deplete LDs in HeLa cells, and miR-122 levels were quantified in control and GW4869-treated HeLa cells, normalized to U6 RNA (n=3). Values with siControl were considered as units. Scale bar 10 μm. F. The effect of SiRNAs against perilipin 2 (siPlin2) on LD number in HeLa cells co-transfected with miR-122 mimic (200nM) (n=10). Data suggest that miRNA levels are lower with fewer LDs. G-H. Effect of expression GFP-Perilipin 2 on LDs (G) and level of cellular miRNAs (H) in HeLA cells transfected with GFP-Perilipin or PCI-neo along with 200nM miR-122 mimic. RNA values quantified by qRT-PCR were normalized to U6 RNA (n=3). Values with pCIneo control were considered as units. Scale bar 10 μm. I. Effect of GFP-Perilin2 expression on LD number determined in HeLa cells having LD with a diameter of more than 100nM diameter. More than 10 cells were analyzed in each case. J-K. Effect of the ATGL inhibitor Atglistatin or ATGLi (100 nM, 8h) on lipid droplets (K) and cellular miRNA content (L) in HeLa cells transfected with 200 nM miR-122 mimic. RNA was quantified by qRT-PCR, and relative levels were normalized to U6 RNA. Scale bar, 10 μm. L. Effect of ATGL inhibitor on LD numbers in HeLa cells. Cells were treated with 100nM ATGLi Atglistatin for 8 h. Data are presented as SEM ± SD, and P values are reported in the respective panels tested for statistical significance. P-values were calculated by a two-tailed paired t-test in most of the experiments unless mentioned otherwise. The fold change of miRNA was calculated by 2^−ΔCt^ method.

However, the importance of LDs in miRNA buffering remains to be verified. The role of lipid droplets in regulating miRNA retention has been examined in cells treated with siPlin2 (On-Target Plus, Dharmacon), a siRNA mix targeting Perilipin2, a key component of LDs. In SiPlin2-treated cells, the number and size of LDs decrease, with an effect on miR-122 content in both control and GW4869-treated conditions, suggesting a possible miRNA storage role for LDs (**Fig.3D-F**). Interestingly, expression of perilipi2 as a GFP fusion construct increased the number and size of LDs, as well as cellular miR-122 content (**Fig.3G-I**). Aglistation or ATGLi, an inhibitor of ATGL that breaks down lipids into fatty acids and thus an inducer of LD formation, increases LD number(44). Treatment of cells with ATGLi increases LD number and increases miRNA-122 content, suggesting a strong correlation between LD number and miRNA content in HeLa cells (**Fig. 3 J-L**).

Lipid droplet association of miRNA also occurs in hepatic cells. We observed increased lipid droplet formation in hepatic cells exposed to high lipid concentrations in the culture medium(30). Cells treated with BSA-Palmitate (BSA-PA)(45) showed abundant lipid droplets in hepatic Huh7 cells, with strong colocalization of miRNA when observed under a confocal microscope (**Fig.4A**). We used biochemical isolation methods to purify LDs enriched for the marker protein ADAR and free of cytosolic and membrane contamination, such as Calnexin(46). To analyze the miRNA content of lipid droplets from control and 300 μm 16h of BSA-PA-treated Huh7 cells. We found a significant association of miR-122 with LDs, and that its abundance increases in LDs isolated from BSA-PA-treated Huh7 cells (**Fig. 4B and C**). However, we failed to detect Ago proteins in the LD-enriched fraction, suggesting Ago2-free miRNAs accumulate on LDs. ATGLi, an inhibitor of Adipose Triglyceride Lipases (ATGL)-a rate-limiting enzyme essential for the breakdown of triglycerides, causes accumulation of LD(47). A similar rise in miR-122 content was also noted in LDs isolated from ATGLi-treated Huh7 cells (**Fig. 4D**). miRNA enrichment with LD is selective, and among the miRNAs tested for enhanced association with LD, let-7a and miR-122 showed significant changes in their association with LD, whereas miR-21 and miR-16 did not change their association with LD in the presence of BSA-palmitate (**Fig. 4E**).

**Figure 4.**
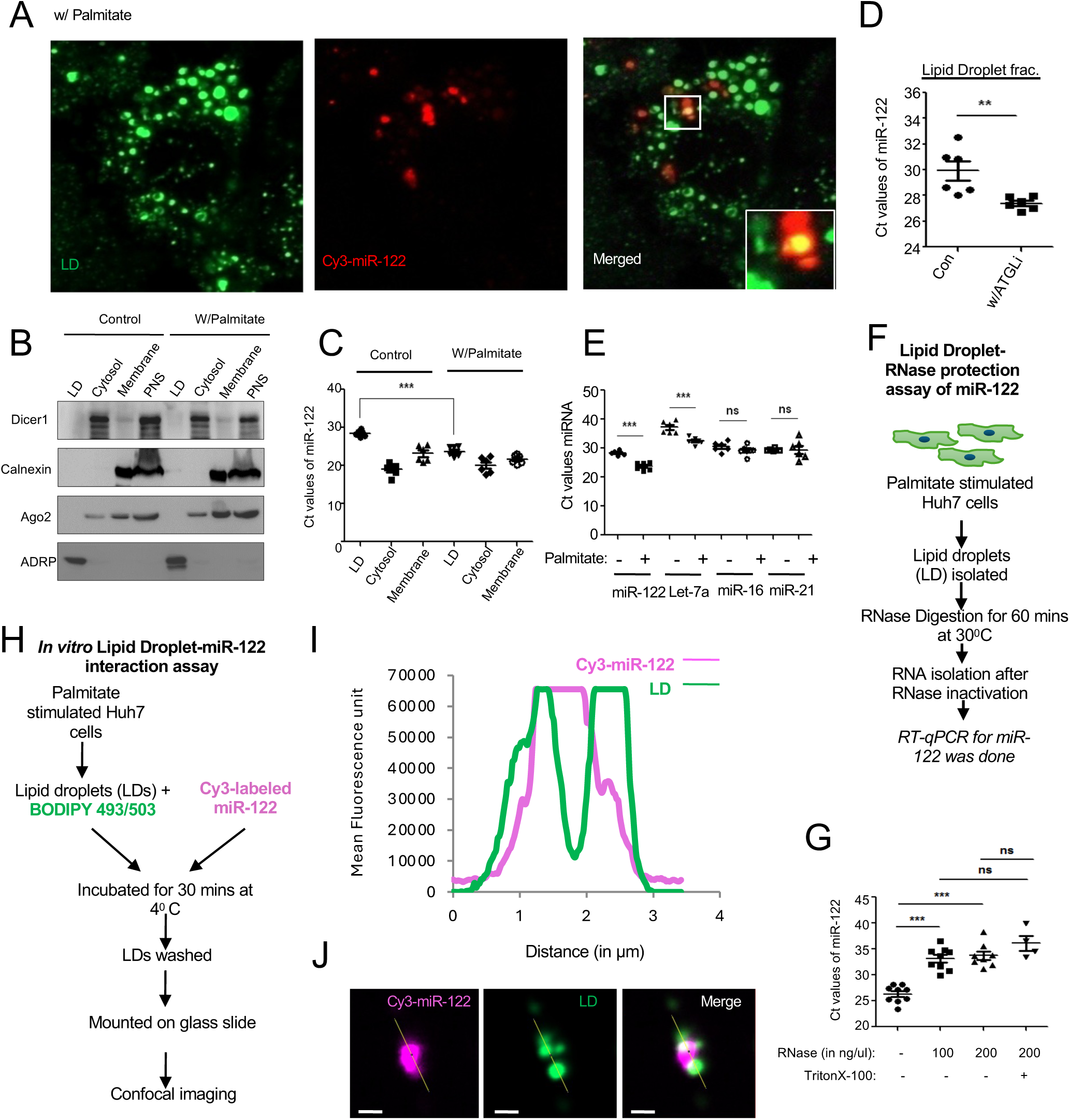
Lipid droplets adsorb Ago2 uncoupled miRNAs. A. Lipid droplets stained with BODIPY 493/503 dye (green) and Cy3-labeled miR-122 (red) are visualized in palmitate-treated hepatic Huh7 cells, showing miRNA and LD colocalization in the zoomed inset. Scale Bar 10 μm. B. Enrichment of Adipose Differentiation-Related Protein (ADRP) in isolated lipid droplets from control and Palmitate-treated Huh7 cells. The Western blot analysis of Calnexin and Ago2 confirms their absence in LD isolated by density gradient centrifugation. C. The miRNA-122 content in different fractions of the control and palmitate-treated Huh7 cells. Showing enhanced miR-122 in LDs isolated from palmitate-treated HuH7 cells. The low Ct value of the miR-122 signifies lower miR-122 present. Value normalized against ADRP levels. D. Enrichment of miR-122 in LDs isolated from ATGLi-treated cells. Control untreated and ATGLi-treated Huh7 cells were used for LD isolation and quantification of miR-122, indicating higher miR-122 in LDs after ATGLi treatment for 8h at 100nM. The low Ct value of the miR-122 signifies lower miR-122 present. E. Relative amount of miRNAs in LD in control and Palmitate-treated Huh7 cells. Relative quantitative estimation by qRT-PCR indicates changes in LD miR-122 and let-7a levels, with no change observed for miR-21 and miR-16. F. Schematic of the experiment with isolated LDs after RNase treatment. Isolated LDs after RNase treatment were used for miRNA estimation done by qRT-PCR and normalized to ADRP. G. Estimation of miR-122 remaining in LD post RNase treatment in the absence and presence of 0.1% Triton X-100 detergent. H. Scheme of *in vitro* LD miR-122 interaction assay. LDs isolated from palmitate-treated Huh7 cells were incubated with ss Cy3-labelled miR-122 before the LDs were washed and reisolated post-interaction with Cy-3 miR-122 (100nM) and analyzed microscopically to find Cy3-miR122 (red) and BODIPY 493/503 dye (green) labelled LD. I. The intensity profile plots for LD and miR-122 over the region of interest defined in panel J. J. The miR-122 and LD signals overlap with those of LDs imaged by confocal microscopy *in vitro*. Scale bar 2 μm. Data are presented as SEM ± SD. * Denotes P value <0.05; ** denotes <0.01 and *** denotes 0<0.001; ns non-significant. The positions of the molecular weight markers are shown in the Western blot panel. P-values were calculated by a two-tailed paired t-test in most of the experiments unless mentioned otherwise. The fold change of miRNA was calculated by 2^−ΔCt^ method.

We subsequently confirmed the membrane association of miRNAs with LDs. With isolated LDs, we performed RNase treatment with or without detergent. After re-isolation of the LDs, followed by RNase treatment, we found a gradual loss of miRNAs with increasing RNase concentration, with a non-significant effect of Triton X-100, suggesting that most miRNAs associated with LDs are present on their surface and don’t need detergent to solubilize them to get them degraded by RNase (**Fig. 4F-G).** To confirm the LD association of the miRNA, *in vitro* assay with isolated LD and Cy3-labeled single-stranded miR-122, was done in the presence of lipid-binding BODIPY dye. After the reaction, the LDs were washed, recovered, and imaged. We found that the RNA and LD signals overlap, suggesting that Cy3-miR-122 is getting associated with LDs after in vitro incubation (**Fig. 4H-J**).

### LD uncouples and stores miRNAs from Ago miRNPs

How do LDs capture single-stranded miRNAs without Ago2 on their surface? This question was addressed in experiments that tested the localization of Ago2 and LD in control versus palmitate-treated Huh7 cells. We observed a proximal localization of Ago2 with LD in palmitate-treated cells, with interaction between LDs and GFP-Ago2 in live-cell imaging experiments (**Fig. S3A-B**), where BSA-palmitate treatment enhances the interaction between LDs and Ago proteins as observed in time-lapse imaging. In experiments with isolated LD and Ago2 miRNPs, immobilized on an HA-Affinity matrix after being isolated from HEK293 cells, we observed a time-dependent loss of miR-122 with Ago2 upon incubation with LDs, suggesting that LDs can decouple miR-122 from Ago2 (**Fig. S3C-D**).

The importance of Ago2 in buffering miRNA levels was further explored in HeLa cells expressing FH-Ago2 (FLAG-HA dual-tagged Ago2), along with a miR-122 mimic, and no major effect of FHA-Ago2 expression on LD number was observed compared with pCIneo control plasmid-transfected cells (**Fig. S4A and B**). However, we observed increased miR-122 levels in FHA-Ago2-expressing cells, suggesting that FHA-Ago2 enhances miRNA stability and function (**Fig. S4C and D**).

### A bipartite role of HuR in miRNA storage in mammalian cells

HuR is a miRNA-binding protein previously shown to be involved in uncoupling mRNA from Ago proteins(12). We have found that HuR depletion negatively affects miRNA storage in LDs. HuR depletion decreases LD number **(Fig. 5A-C)** and reduces miRNA levels in both control cells and GW4869-treated cells **(Fig. 5D).** The reduction in miR-122 levels in HuR-depleted cells is independent of Bafilomycin, as Bafilomycin cannot affect the reduction of miRNAs in HuR-depleted cells (**Fig. 5E**). This observation strongly suggests that LD compartmentalization of miRNAs requires uncoupling from Ago proteins, a process facilitated by Hu(48). To confirm the role of HuR in controlling LD association of miRNAs, we performed *in vitro* LD miRNA-loading experiments using isolated LDs and Ago2 miRNP in the presence and absence of recombinant HuR protein, known to bind miR-122(12). We found a HuR-dependent enhancement of miR-122 association with LDs in the presence of recombinant HuR protein, suggesting a direct role of HuR in uncoupling Ago2 from miRNAs and in subsequent LD binding of single-stranded miR-122 (**Fig. 5F-H**).

**Figure 5:**
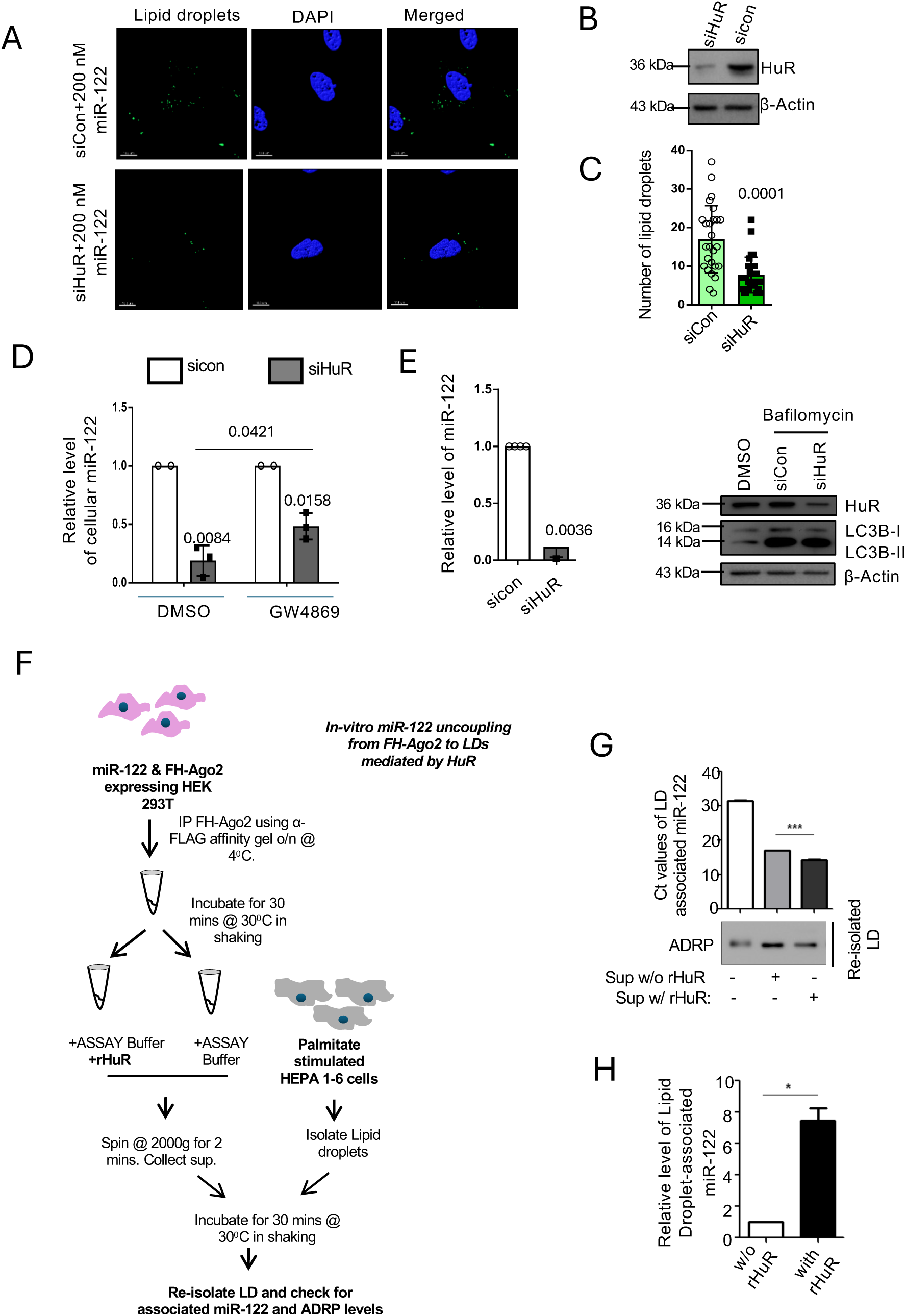
Effect of HuR depletion on the lipid droplets and miRNA levels in HeLa cells. A-C. Treatment of siCon and siHuR (20nM, 72h) on lipid droplets in HeLa cells (A). LDs were visualized using the BODIPY 493/503 dye (green) in siCon and siHuR-transfected cells. Scale bar, 10 μm. The western blot confirms HuR depletion. Actin was used as a loading control (B). Lipid droplet numbers were calculated from 25 cells (C). Scale bar, 10 μm. D-E. The effect of siHuR on miRNA levels in HeLa cells transfected with miR-122 in control DMSO and GW4969-treated cells (D). The effect of Bafilomycin on miRNA levels in siHuR-treated cells is shown (E). In HeLa cells, the effect of HuR depletion on LC3BII levels in control and siHuR-treated cells was confirmed by Western blot (E, lower panel). β-Actin was used as a loading control. F-H. Schematic of the experiments performed with isolated LDs incubated with Ago2 miRNPs isolated from HEK293 cells expressing FH-Ago2 and miR-122 (F). Post assay, the reisolated LDs were measured for miR-122 content by qRT-PCR. Recombinant HuR was used at a 200 nM concentration (G). The fold change of miR-122 in LDs in control and rHuR-treated LDs was measured and plotted (H). All data are from three experimental replicates and are presented as SEM ± SD. P values are reported in the respective panels tested for statistical significance. P-values were calculated by a two-tailed paired t-test in most of the experiments unless mentioned otherwise. The fold change of miRNA was calculated by 2^−ΔCt^ method. The positions of the molecular weight markers are shown in the Western blot panel. P values are either shown within the panels or * Denotes P value <0.05; ** and *** denotes 0<0.001.

By uncoupling miRNAs from Ago2 and reversibly binding to miRNAs, HuR also facilitates the entry of miRNAs into the endosomal lumen, enabling subsequent export via EVs in animal cells. HuR is also involved in stress-induced export of miR-122 from hepatic, macrophage, and astroglial cells. Thus, HuR overexpression is expected to have a miRNA-reducing role by promoting miRNA export from cells transfected with an excess miR-122 mimic beyond cellular threshold levels. As the amount of miRNA accumulated inside human cells increases, ectopic expression of HuR reverses miRNA storage, and cellular miR-122 content decreases with HA-HuR expression, a result consistent with observations in hepatic cells, where HuR causes export of miRNA to reduce miRNA content, while EV-associated miR-122 levels increase (**Fig. 6A-D**). Interestingly, HA-HuR decreases lipid droplet number, and miRNA localization was not observed in HA-HuR-expressing HeLa cells transfected with 200 nM of miR-122 mimic (**Fig. 6C, E-F**). These apparent contradictory results could be explained by a possible bipartite role of HuR in miRNA buffering. While HuR facilitates endosomal targeting and export of miRNAs, a process dependent on miRNA binding, it may also require cooperative interactions among miRNA cargoes to facilitate export via EVs(49). Whereas HuR is essential for storing miRNAs in a nonfunctional form in lipid droplets. We have found that with increasing palmitate treatment, miR-122 content accumulates in lipid droplets in hepatic cells, and this accumulation is associated with increased HuR levels(30) (**Fig. 4A**; **Fig. 6G**).

**Figure 6.**
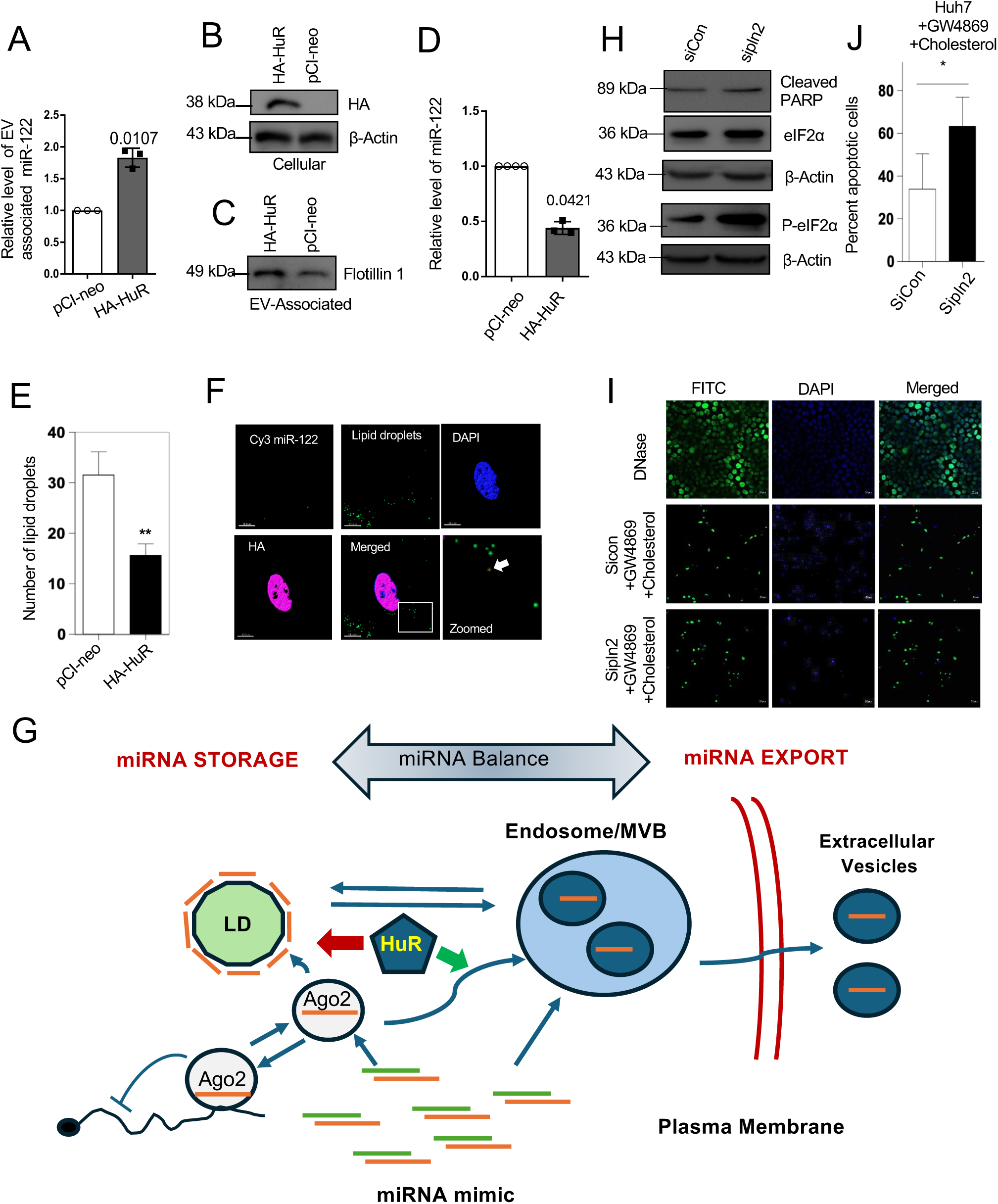
miRNA sensing by HuR is required for miRNA homeostasis. A-D. Effect of HA-HuR expression on EV-associated miR-122 isolated from cells transfected with pCIneo or HA-HuR expression plasmids (A). HA-HuR expression is confirmed by Western blot using an HA-specific antibody (B). A Western blot for Flotillin-1 was used to normalize EV content for quantitative measurement of miR-122 in EVs. The change in cellular miR-122 content was measured by qRT-PCR (D) using U6 RNA as a control. E. Effect of HA-HuR expression on lipid droplet number in HeLa cells co-transfected with 200 nM miR-122 mimic. The change in the number of LD has been quantified in HeLa cells (n=20). F. Failed localization of miR-122 to LD in cells expressing HA-HuR. Cell nucleus stained with DAPI and Cy-3 miR122 was detected in red. LD was visualized with BODIPY dye (green) while HA-HuR was in Cyan. Scale bar, 10 μm. G. A working model of how miRNA homeostasis is regulated in human cells by lipid droplet storage and extracellular export via endosomal pathways. Both pathways are controlled by HuR, which facilitates Ago2-miRNA unloading to favor miRNA storage and endosomal entry. HuR-miRNA complexes may be accessible to LDs and endosomes, and HuR-interacting proteins such as STX5 or RalA, as well as HuR modifications such as ubiquitination, favor the endosomal/exosome-mediated export. However, in cells compromised for export, LD-mediated miRNA storage may affect miRNA function and protect cells from miRNA toxicity. H-J. Effect of PB depletion on apoptotic death of Huh7 cells exposed to MCD-cholesterol concentrate for 8 h and pre-treated with GW4869 for 24h. The failure to store miR-122 alters the expression of cleaved PARP and phosphorylated eIF2α, stress markers (H). The TUNEL assay also shows a higher number of apoptotic cells lacking LDs following siPlin2 treatment (I). Quantitative changes in apoptotic cell numbers are calculated and plotted in panel J. All data are from three experimental replicates and are presented as SEM ± SD. P-values were calculated by a two-tailed paired t-test in most of the experiments unless mentioned otherwise. The fold change of miRNA was calculated by 2^−ΔCt^ method. The positions of the molecular weight markers are shown in the Western blot panel. P values are either shown within the panels or * Denotes P value <0.05; ** and *** denotes 0<0.001.

### miRNA storage in lipid droplets prevents apoptosis in hepatocytes exposed to high lipids

Our previous research demonstrated that excess miR-122 activity under starvation-induced stress or high-lipid conditions induces apoptotic cell death in hepatic cells(12, 30). This is linked to the accumulation of miR-122 in cells treated with GW4869(12). Therefore, reducing or buffering miR-122 activity is crucial for regulating miRNA activity and may influence downstream pathways related to apoptosis. We propose that lipid droplets serve as ideal storage sites for “inactive” miRNAs in mammalian cells, and accumulating miRNAs in lipid droplets acts as a safeguard against miRNA toxicity in hepatic cells exposed to high miR-122 levels due to high lipids. Our experiments showed increased apoptosis in miR-122-loaded hepatocytes treated with Perilipin 2-specific siRNAs compared to non-targeting siRNA control, evidenced by increased PARP cleavage and eIF-2α phosphorylation in GW4869-treated hepatic cells exposed to high cholesterol(30). These findings highlight the importance of lipid droplets in regulating the availability of cellular miR-122, necessary for preventing apoptosis in lipid-challenged hepatic cells (**Fig. 6H-J**). Interestingly, ATGLi treatment, which induces LDs and storage of miRNA on LD (**Fig. 3J-L**), reduces the phosphorylated eIF2a level in cholesterol-treated Huh7 cells, reconfirming LDs’ role in controlling the lipotoxic stress response and possibly affecting miR-122 activity in hepatic cells (**Fig. S5**).

Overall, this data suggests that miRNA levels are buffered in mammalian cells, with HuR protein serving as a key dual regulator in this process. As a miR-122 sensor, HuR first disconnects miRNAs from Ago-miRNPs, promoting their export via extracellular vesicles in cells overloaded with miRNAs. Additionally, HuR is crucial for miRNA-association with LDs, a process facilitated by Ago2-mediated release of miRNAs. Although HuR’s role in miRNA export may be redundant—possibly mediated by other proteins such as STX5(13)—it remains necessary for miRNA targeting to LDs (**Fig. 6G**).

## Discussion

How could miRNA activities be regulated in mammalian cells? Recent work from our group has revealed the mitochondrial storage function of subsets of miRNAs targeted to mitochondria, which augment the stress response(50). The role of the RNA processing body (P-body) in mRNA and miRNA storage has been described as a mechanism for controlling miRNA activity under stress conditions(16). The importance of miRNA export as a mediator of intercellular exchange of epigenetic information has been explored, and its role in exporting specific miRNAs to balance miRNA activity has been shown in hepatic cells under stress(12) and in immune cells under specific needs(48). miRNAs are packaged into EVs in sequence and context-dependent manner, regulated by several RNA-binding proteins that facilitate miRNA export from mammalian cells(12, 49, 51, 52), allowing effective communication with adjacent cells and tissues. Suggesting that miRNA export is selective and specific.

The proposed existence of miRNA-sensing RNA-binding proteins (RBPs) explains the fine-tuning of miRNA activity in mammalian cells(53). RBPs bind specific sequences or structures in pri-miRNA and pre-miRNA to promote or inhibit their maturation or even target recognition(54). Proteins such as hnRNPA1 and KSRP recognize terminal loop structures, inducing conformational changes that promote cleavage by Drosha-DGCR(55). Proteins such as QKI and HuR stabilize mature miRNAs, extending their half-life(56, 57). Exonucleases like XRN1/2 and PNPase degrade miRNAs, enabling rapid turnover(58, 59). RBPs like HuR or DND1 can also bind near target sites to mask them, preventing RISC repression(60, 61). Some RBPs remodel mRNA secondary structures, exposing target sites to miRNA-RISC. This fine-tuning can also be achieved by miRNA sponges, which are naturally non-coding RNAs expressed under specific conditions and thereby deactivate miRNA function by binding to miRNAs, the 22-nt-long regulators of gene expression(62). The question of how miRNA export wins over other miRNA-degradation or associated pathways remains unresolved. Several predicted mechanisms exist in which specific post-transcriptional modifications of miRNAs initiate degradation or export(63). Specific post-transcriptional modifications that initiate the degradation of mature microRNAs (miRNAs) include uridylation (addition of uridine nucleotides) and, in certain contexts, adenylation (addition of adenine nucleotides)(64). A distinct mechanism involving high-complementarity target RNAs (Target-Directed miRNA Degradation, or TDMD) also triggers decay by exposing the miRNA 3’ end for enzymatic attack. Recent work has defined adenylation of miRNAs coupled to degradation or export. The turnover and export of miRNAs have also been shown to depend on the availability of target mRNAs, while the impact of target mRNAs can be positive or negative, as published evidence suggests both possibilities(15, 65).

Our data suggest the existence of a buffering system, initiated by sensing miRNA content via the miRNA-binding protein HuR, that exports Ago2-uncoupled miRNAs to balance miRNA turnover as and when required. With increasing concentrations of miRNA-122 mimic used for transfection of HeLa cells, we observed a decrease in cellular miRNA content after the cells reached a threshold, and blocking miRNA export recovered miRNA levels. Cellular miRNA levels increased with increasing miRNA mimic used for transfection. The excess miRNA, which remains non-functional, is stored in lipid droplets. Why is there a drop in miRNA levels after the cellular miRNA concentration reaches a threshold? Possibly because Ago proteins become saturated, leaving excess miRNAs targeted for export, which may initiate a mechanism of miRNA modification that allows a synergistic effect on export of other miRNAs(49), as endogenous miRNA let-7a levels decrease with excess miR-122 present in the system at 200 nM miR-122 used. The miRNAs that accumulate in LDs may serve as a safeguard mechanism that allows cells to adapt when miRNA demand increases (e.g., when miR-122 activity is required to reverse lipotoxic stress). The LD-associated miRNAs are single-stranded and may not require Dicer1 for reloading onto Ago2. This has been observed with EV-internalized miRNAs that get loaded onto recipient cells’ Ago2 in a Dicer1-independent manner(66). The reversible storage of miRNAs in lipid droplets raises questions, such as how charged molecules, including miRNAs, remain attached to the lipid membrane. There has been a report of RNA binding to the lipid membrane via the glycosidic moiety, resulting from RNA glycosylation(67). Lipids on the LD may interact with miRNAs in a similar manner. Alternatively, HuR-bound miRNAs may be a substrate for LD-associated proteins, such as Perilipin 2, which, by interacting with HuR, facilitates miRNA binding to LDs. Perilipin 2 (Plin2) is a major protein associated with the surface of lipid droplets and plays a crucial role in maintaining LD structure and regulating neutral lipid metabolism(68), and may be involved in RNA interaction. Perilipin 2 (PLIN2) interacts with various RNA-binding proteins (RBPs), especially Rab18, which binds PLIN2 and ACSL3 to regulate lipid droplet behavior(69, 70). Additionally, it interacts with proteins like ABHD5, DESI2, HTATIP2, KIF1B, POGZ, SFT2D2, and WFS1, many of which have RBP roles or participate in RBP-related pathways. These interactions influence lipid droplet dynamics, gene expression, and overall cellular function, thereby linking lipid metabolism to RNA processing(70, 71). These crucial questions are essential for understanding how miRNAs are adsorbed onto lipid droplets. The factors that regulate the balance between lipid droplet storage and miRNA export, and vice versa, are essential for identifying candidates to manipulate the process. We posit that HuR plays an essential role in both augmenting miRNA export and miRNA LD loading. It is possible that dependence on HuR ubiquitination, which allows its unbinding from miRNAs on endosomes(12), and the availability of STX5, a miRNA-interacting protein(13), can determine which pathways HuR adopts to regulate miRNA fate. Overall, this investigation has revealed a new role for lipid droplets in regulating gene expression in mammalian cells, including hepatic cells, by modulating miRNA activity to protect cells from apoptotic death.

## Experimental Procedures

### Cell culture, cell transfection, chemicals, and plasmids

Human HeLa, HEK293, and Huh7 cells were cultured in Dulbecco’s Modified Eagle’s Medium (DMEM; Gibco-BRL), supplemented with 2mM L-glutamine, 10% heat inactivated fetal bovine serum (FBS), and 1% penstrep (Gibco). Plasmid transfections were done using Lipofectamine 2000 (Invitrogen), following manufacturer’s protocol. The EV export blocker GW4869 was purchased from Calbiochem CA and was used at 10μM concentration for 24h (12). Atglistatin (Sigma) was applied at a final concentration of 20 μM for 16 h. Bafilomycin (Sigma) was applied at a final concentration of 100nM for 8 h. Plasmid F-HA-Ago2 was kind gift from Tom Tuschl [previously described in(12)]. Plasmids NHA-GW182b and GFP-Dcp1a were kind gifts from Witold Filipowicz [previously described in (18)]. Plasmid HA-HuR was a kind gift from Witold Filipowicz (previously described by Mukherjee et al., 2016 [9]). Plasmids RL-con and RL-3xb-let-7a and RL-3xb-miR-122 were kind gifts from Witold Filipowicz [previously described in (16)]. Plasmid GFP-perilipin 2 was purchased from Origene. Plasmids FF and pci-Neo were purchased from Promega.

### siRNA and miRNA mimic transfections

siRNA and miRNA mimic transfections were performed in RNAimax (Invitrogen), following the manufacturer’s protocol. 20 picomoles of siRNA was used per well of a 24-well plate. siControl, siHuR (human), and si-perilipin 2 (human) were purchased from Dharmacon, and mirVana miRNA hsa-miR-122-5p, hsa-miR-155-5p, and hsa-miR-146a-5p mimics were purchased from Ambion (Life technologies).

### Western Blotting

Protein samples were heated in 1X SDS dye (5X SDS dye containing 312.5 mM Tris-HCl pH 6.8, 10% SDS, 50% glycerol, 250 mM DTT, 0.5% bromophenol blue, and 10% β-mercaptoethanol) for 10 minutes at 95 ⁰C. Subsequently, SDS-PAGE was performed. The proteins were then transferred to a PVDF nylon membrane (activated with methanol), followed by blocking in TBS with 0.1% Tween-20 and 3% BSA (1X TBST with BSA). After blocking, the membrane was incubated overnight at 4 ⁰C with primary antibody in 3% BSA in 1X TBST. The membrane was then washed three times for 5 minutes each with 1X TBST at room temperature, and secondary antibodies conjugated with Horseradish peroxidase (dilution 1:8000) in 3% BSA in 1X TBST were added for 1 hour at room temperature. Post-incubation, the membrane was washed again thrice with 1X TBST for 5 minutes each. The antigen-antibody complex was visualized using West Pico Chemiluminescent or West Femto Maximum Sensitivity substrates following the manufacturer’s instructions (Thermo Scientific). Imaging was performed with a UVP BioImager 600 system equipped with VisionWorks Life Science software (UVP) V6.80. Antibody dilutions used were: flotillin 1 (1:500), HA (1:1000), HuR (1:1000), LC3B (1:1000), and β-actin (1:10000).

### BSA-Palmitate and cholesterol treatment

A 0.2 M ethanolic solution of palmitic acid (Sigma) was conjugated to fatty acid-free BSA Fraction V (Sigma) in KRBH buffer, which contains 3.6 mM KCl, 135 mM NaCl, 10 mM HEPES, 5 mM NaHCO3, 0.5 mM NaH2PO4, 0.5 mM MgCl2, 1.5 mM CaCl2, 3.5 mM D-glucose, and 21% BSA. As a control, an equal volume of ethanol was added to KRBH buffer. The solutions were shaken at 37°C for 6 hours and then filter-sterilized. The resulting stock of BSA-palmitate had a final concentration of 10 mM with a BSA: palmitic acid ratio of 1:3. Both BSA-palmitate and BSA-only control solutions were stored at −20°C and pre-warmed at 37°C for 15 minutes before being applied to 70% confluent Huh7 or Hepa 1-6 cells in fresh DMEM complete media. Treatments lasted 16 hours. Methyl-β-cyclodextrin–conjugated cholesterol obtained from Gibco (catalog no.: 12531-018) was added from a 250× stock to Huh7 cells in culture at a final concentration of 1x for 8 h (30 mg/l effective cholesterol in 1×)(30).

### RNA isolation and real time-PCR

RNA was isolated using TriZol reagent (Invitrogen) following the manufacturer’s protocol. Real-time analysis of miRNA via a two-step RT-PCR was performed with TaqMan chemistry (Applied Biosystems) on a Bio-Rad CFX96TM real-time system. For cellular samples, 200 ng of RNA was used, while an equivalent volume was used for EV samples. U6 snRNA served as the endogenous control for cellular samples. Quantification employed specific primers for human miR-122, miR-155, miR-146a, let-7a, miR-21, and U6 snRNA. The CT values of target miRNAs were normalized against U6 snRNA CT values to determine relative miRNA levels. EV samples were normalized by band intensity of Flotillin-1 using ImageJ software.

mRNA levels were estimated using SYBR Green-based real-time PCR on a 7500 Applied Biosystems system. For cellular samples, 200 ng of RNA was used. Specific primers targeted human CAT1, Aldolase A, and GAPDH, with GAPDH serving as the endogenous control. The CT values of target mRNAs were normalized to GAPDH CT values, and relative mRNA levels were calculated.

### EV isolation and characterization

Plasmid transfection was performed in a 12-well plate. The next day, cells from one well were transferred to a well of a 6-well plate, followed by addition of EV-depleted media (prepared from EV-depleted FBS). The cells were then cultured for 24-48 hours until reaching 70-80% confluency, after which EVs were isolated. The cell supernatant was centrifuged at 2000g for 15 minutes at 4°C to remove cell debris, then spun again at 10000g for 30 minutes at 4°C. The supernatant was filtered through a 0.22μm syringe filter, and EVs were pelleted by two rounds of ultracentrifugation at 31200rpm for 90 minutes at 4°C on a sucrose cushion. The EV pellet was resuspended in either 1X PBS or 1X PLB for subsequent use.

### Luciferase Assay

To measure miRNA-mediated repression, a dual-luciferase assay was conducted. Cells seeded in 12-well plates were transfected with 50 ng of Renilla Luciferase (RL) constructs—either containing three let-7a binding sites at the 3’-UTR (RL-3xB-let-7a) or no let-7a binding sites (RL-Con). They were co-transfected with a Firefly Luciferase plasmid (FF) for normalization. The cells were lysed using 1x Passive Lysis Buffer (Promega), and luciferase activity was measured with the Dual-Luciferase Reporter Assay kit (Promega). Readings were recorded on a VICTOR X3 Plate Reader (PerkinElmer, Waltham, MA). For each sample, three technical and three biological replicates were included. Fold repression was calculated as the ratio of Firefly-normalized RL-Con expression to RL-3xB-let-7a expression as described above.

### Lipid Droplet Isolation

Lipid droplets (LDs) were isolated using a floatation gradient ultracentrifugation method. Approximately 3 × 10^7 Huh7 cells, either stimulated with or without palmitate, were homogenized in Buffer A (isotonic, containing 20 mM Tricine, 250 mM sucrose, 1x Protease inhibitor (Roche), 40 U/ml RNase inhibitor (Applied Biosystems), pH 7.8). The homogenate was clarified by centrifugation at 1000 x g for 5 minutes at 4 °C to produce about 3.5 ml of post-nuclear supernatant (PNS), which was loaded into an ultracentrifuge tube. Then, 0.5 ml of Buffer B (containing 20 mM HEPES, 100 mM KCl, 1x Protease inhibitor, 40 U/ml RNase inhibitor, pH 7.4) was carefully layered on top of the PNS, creating a discontinuous gradient. This was ultracentrifuged at 1,00,000 x g for 1 hour and 15 minutes at 4°C. After centrifugation, the LDs appeared as a white, milky layer floating on top of the tube. They were gently collected from the top. To remove organellar contaminants, the LDs were washed three times in Buffer B by centrifugation at 20,000 x g for 5 minutes at 4°C. The cytosolic fraction was collected from the middle of the tube, and the total membrane fraction was obtained as a pellet.

### In vitro assays with isolated Lipid Droplets

The in vitro Ago2-Dicer1 interaction assay was conducted using isolated LDs from palmitate-stimulated Huh7 cells. NH-Dicer1 transfected HEK293 cells were homogenized in Buffer A with a Dounce homogenizer. The homogenate was then clarified by centrifugation at 1000 x g for 5 minutes to obtain the post-nuclear supernatant (PNS). Meanwhile, LDs were isolated from palmitate-treated Huh7 cells and dissolved in Buffer B. The HEK293 PNS was incubated with the isolated LDs under shaking conditions for 60 minutes at 37°C. As a control, HEK293 PNS was incubated with an equal volume of Buffer B. After the reaction, NH-Dicer1 was immunoprecipitated from the mixture, and the associated Ago2 was detected via western blot analysis. For the in vitro RNase assay of LD-associated miR-122, LDs were first isolated from around 3 × 10^7 Huh7 cells that had been stimulated with palmitate. The isolated LDs underwent RNase A digestion at concentrations of 100 and 200 ng/µl at 300C for 60 minutes, with or without 1% Triton X-100. Afterwards, RNase A was inactivated, and RNA was extracted from the reaction mixture using Trizol LS reagent (Invitrogen). Finally, miR-122 levels were quantified via RT-PCR from the remaining RNA. The dissociation of miR-122 from Ago2 in the presence of LDs was examined using an in vitro Ago2-miR-122 uncoupling assay with LDs. HEK293 lysates expressing FH-Ago2 and miR-122 were homogenized in Buffer A, and PNS was collected by centrifugation at 1000 x g for 5 minutes at 4°C. The post-nuclear supernatant was then incubated with LDs isolated from palmitate-stimulated Huh7 cells for 15, 30, and 60 minutes at 37°C. FH-Ago2 was subsequently immunoprecipitated. RNA was extracted from half of the immunoprecipitated sample using Trizol (Invitrogen), and miR-122 levels were measured by RT-PCR. The remaining half was analyzed via western blot to confirm the amount of FH-Ago2 immunoprecipitated.

### TUNEL Assay

TUNEL assay used Click-iT™ TUNEL Alexa Fluor 594 kit (Invitrogen). Huh7 cells expressing NH-DICER1 were grown on 12-well coverslips, then treated with cholesterol for specified times. After treatment, cells were fixed with 4% paraformaldehyde in PBS for 30 mins and permeabilized with 0.25% Triton X-100 for 20 mins. The TUNEL assay was then performed to incorporate modified dUTPs using terminal deoxynucleotidyl transferase, followed by a click reaction for fluorescent detection. Coverslips were counterstained for NH-DICER1 with anti-HA primary antibodies and Alexa fluor-488 secondary antibodies (Invitrogen). Quantification involved calculating the percentage of green (NH-DICER1 positive) and non-green cells positive for TUNEL (red).

### Confocal microscopy and post-capture image analysis

Cells on gelatin-coated coverslips were transfected and treated as described, then fixed with 4% paraformaldehyde in PBS for 30 mins at room temperature in the dark. For immunofluorescence, fixed cells were permeabilized and blocked at room temperature for 30 mins with 3% BSA in PBS containing 0.1% Triton X-100, then incubated overnight at 40°C with primary antibodies. Non-specific binding was washed away with PBS three times (5 mins each). Cells were stained with fluorochrome-tagged Alexa fluor secondary antibodies (Invitrogen) for 1 hour at room temperature, then washed three times with PBS. Coverslips were mounted on slides with VECTASHIELD antifade mounting media. For lipid droplet staining, cells were incubated overnight with BodipyTMC12 558/568 or probed with BodipyTM 493/503 (Invitrogen) diluted in PBS for 30 mins at room temperature. Imaging was performed with a Zeiss LSM800 confocal microscope. All post-capture image processing used Imaris 7. For colocalization, images were thresholded, and Pearson’s coefficient was calculated with the Coloc. plugin. 3D reconstructions of z-stacks used the Surpass plugin. Lipid droplets per cell were counted with the particle generator in Imaris 7.

### Preparation of recombinant HuR

Recombinant HuR protein was purified as previously described(12). HuR was expressed in E. coli using a pET42a(+) construct in BL21DE3 cells. Cultures were induced with IPTG overnight, lysed with a buffer containing Tris–HCl, KCl, MgCl2, β-mercaptoethanol, imidazole, Triton X-100, glycerol, lysozyme, and protease inhibitors, then sonicated. Lysates were clarified and incubated with Ni-NTA agarose beads at 4°C for 4 hours. Beads were washed with a similar buffer on a rotator at 4°C. His-tagged HuR was eluted with buffers of increasing imidazole concentration, each for 15 minutes at 4°C. The purified protein was stored at −80°C. (12, 52).

### Statistical Analysis

Data analysis was performed using GraphPad Prism 5.00 (Graph Pad, San Diego) on data plots from experiments conducted in triplicate unless indicated otherwise. P values were calculated with Student’s t-test, considering results significant at P<0.05. Error bars represent SEM±s.d. (12, 52).

## Data Availability Statement

All relevant data supporting the key findings of this study are available within the article and the Supplementary Information file. No protein or RNA Sequencing data are used in the manuscript for deposit in repository domains.

## Acknowledgements

We thank Witold Filipowicz for providing the various constructs used in this study. We acknowledge the funding from the Department of Science and Technology (DST), Government of India, the Council for Scientific and Industrial Research (CSIR), and the University Grants Commission (UGC) for their support. SNB was supported by the Swarnajayanti Fellowship (DST/SJF/LSA-03/2014-15) from DST, Govt. of India. This work also received support from a High-Risk High Reward Grant (HRR/2016/000093) from DST and the CEFIPRA project grant 6003-J. Currently, SNB is supported by the Start-Up Support Grant from the University of Nebraska, USA, and the Lieberman Research Award, Department of Anesthesiology, UNMC, supporting K.M.

## CRediT Author Contribution

Sreemoyee Chakraborty: Data Curation, Formal Analysis, Investigation, Methodology. Diptankar Bandyopadhyay : Formal Analysis, Investigation, Methodology. Kamalika Mukherjee: Data Curation, Formal Analysis, Investigation, Writing-Review and Editing, Conceptualization, Methodology, Supervision. Suvendra N Bhattacharyya : Data Curation, Formal Analysis, Investigation, Writing-Review and Editing, Conceptualization, Funding Acquisition, Methodology, Supervision, Writing-Review and Editing.

## Supplementary Information

### Supplementary Figures and Legends

**Fig. S1.**
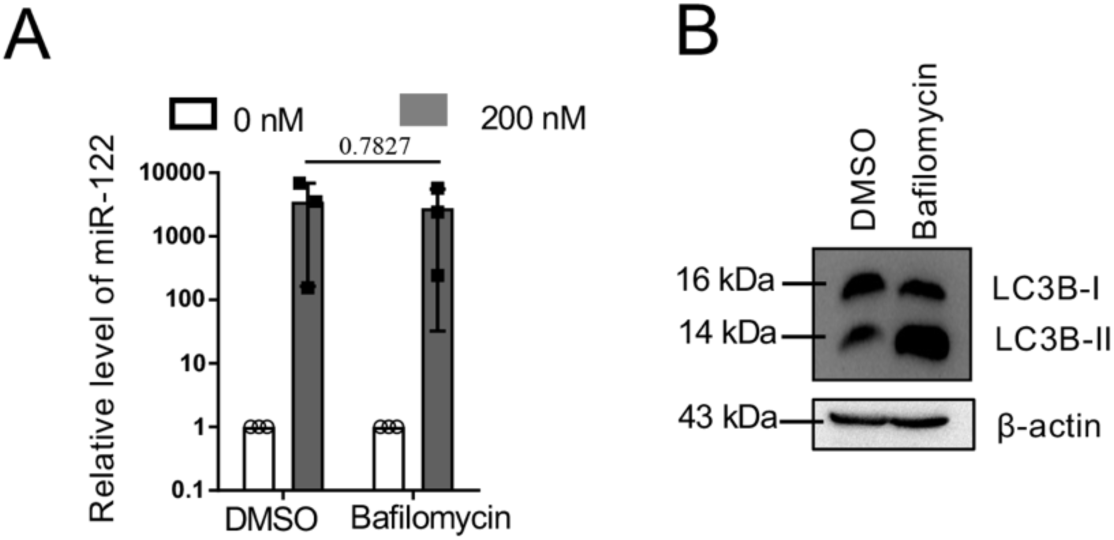
Effect of Bafilomycin on cellular miRNA levels in mammalian cells. A. Relative levels of miR-122 in Bafilomycin (8h) treated HeLa cells transfected with 0 and 200nM miR-122 mimic. Values were normalized to U6 RNA, and results obtained with DMSO and without miR-122 transfection were used as controls for the experiment performed in triplicate. B. Bafilomycin treatment effect on autophagy pathway was confirmed by an increase in the protein levels of LC3B-II (right panel). β-actin served as the loading control for Western blotting. Data are presented as SEM ± SD, and P values are reported in the respective panels tested for statistical significance. The positions of the molecular weight markers are shown in the Western blot panel. P-values were calculated by a two-tailed paired t-test in most of the experiments unless mentioned otherwise. The fold change of miRNA was calculated by 2^−ΔCt^ method.

**Figure S2.**
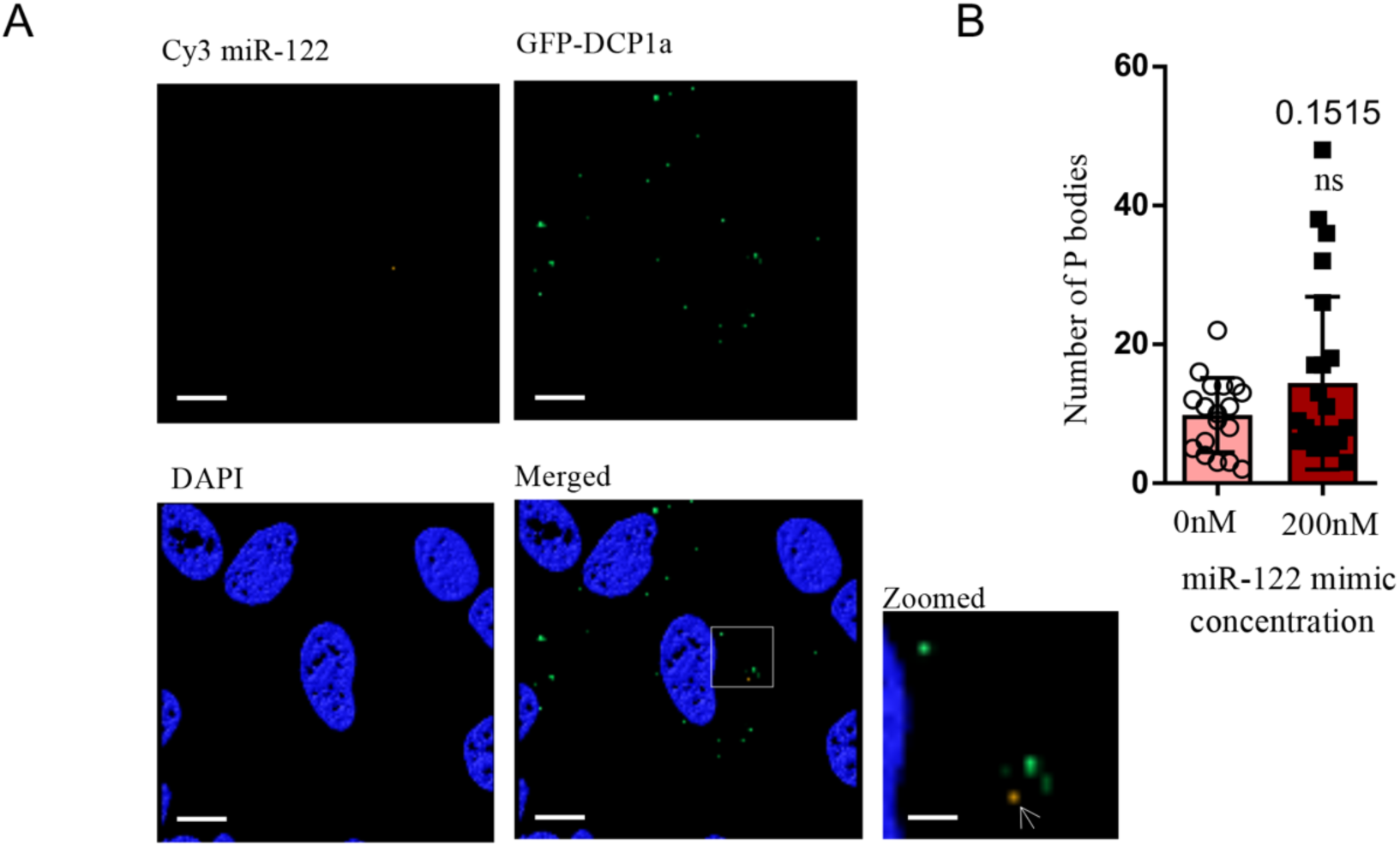
Effect of miR-122 on Dcp1a-positive P-bodies in HeLa cells. A. Localization of Cy-3 miRNA and GFP-Dcp1a in HeLa cells transfected with 200nM Cy-miR-122. Zoomed part showing P-bodies and miR-122 foci. Scale bars are 10μm and 2 μm (zoomed panel). B. Effect of miR-122 mimic transfection on Dcp1a-positive P-bodies number. Visible P-bodies were counted in each case and plotted (n=20). Data are presented as SEM ± SD, and P values are reported in the respective panels tested for statistical significance. P-values were calculated by a two-tailed paired t-test in most of the experiments unless mentioned otherwise.

**Figure S3.**
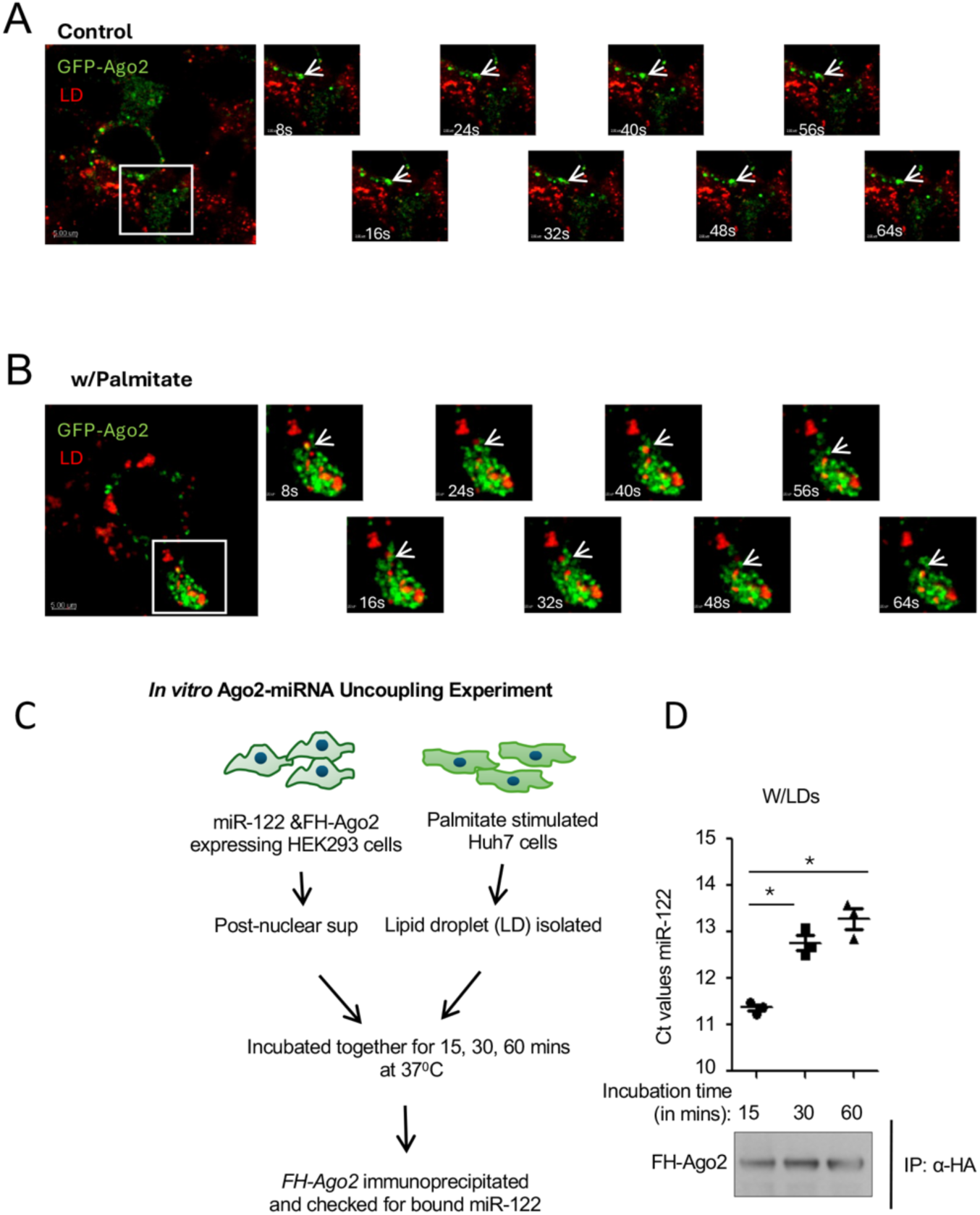
Ago2 Uncoupling of miRNA by Lipid droplets. A-B. Time-lapse imaging of GFP-Ago2-expressing control (A) and BSA-Plamitate (B) treated Huh7 cells. GFP-Ago2 is shown in green, and LD is stained with Red/Deep red BODIPY were imaged at 10-second intervals to show the interaction of LD with Ago2. Confocal super-resolution imaging was performed. Scale bar, 10 μm. Arrowhead indicating LD proximal to Ago2. C. LD uncouples miRNA from Ago2 in the *in vitro* LD-Ago2 interaction assay. We incubated isolated LDs with FLAG-HA-tagged Ago2 (FH-Ago2) and miR-122 miRNPs (100nM) isolated from HEK293 cells expressing FH-Ago2 and miR-122 (C). After the reaction, FH-Ago2 was reisolated and quantified for miR-122 content. The amount of miRNA in reisolated Ago2 immunoprecipitated was measured over time and quantified by qRT-PCR (D). Value normalized against Ago2 level. Data are presented as SEM ± SD of three experimental replicates. * Denotes P value <0.05. P-values were calculated by a two-tailed paired t-test in most of the experiments unless mentioned otherwise. The fold change of miRNA was calculated by 2^−ΔCt^ method.

**Figure S4.**
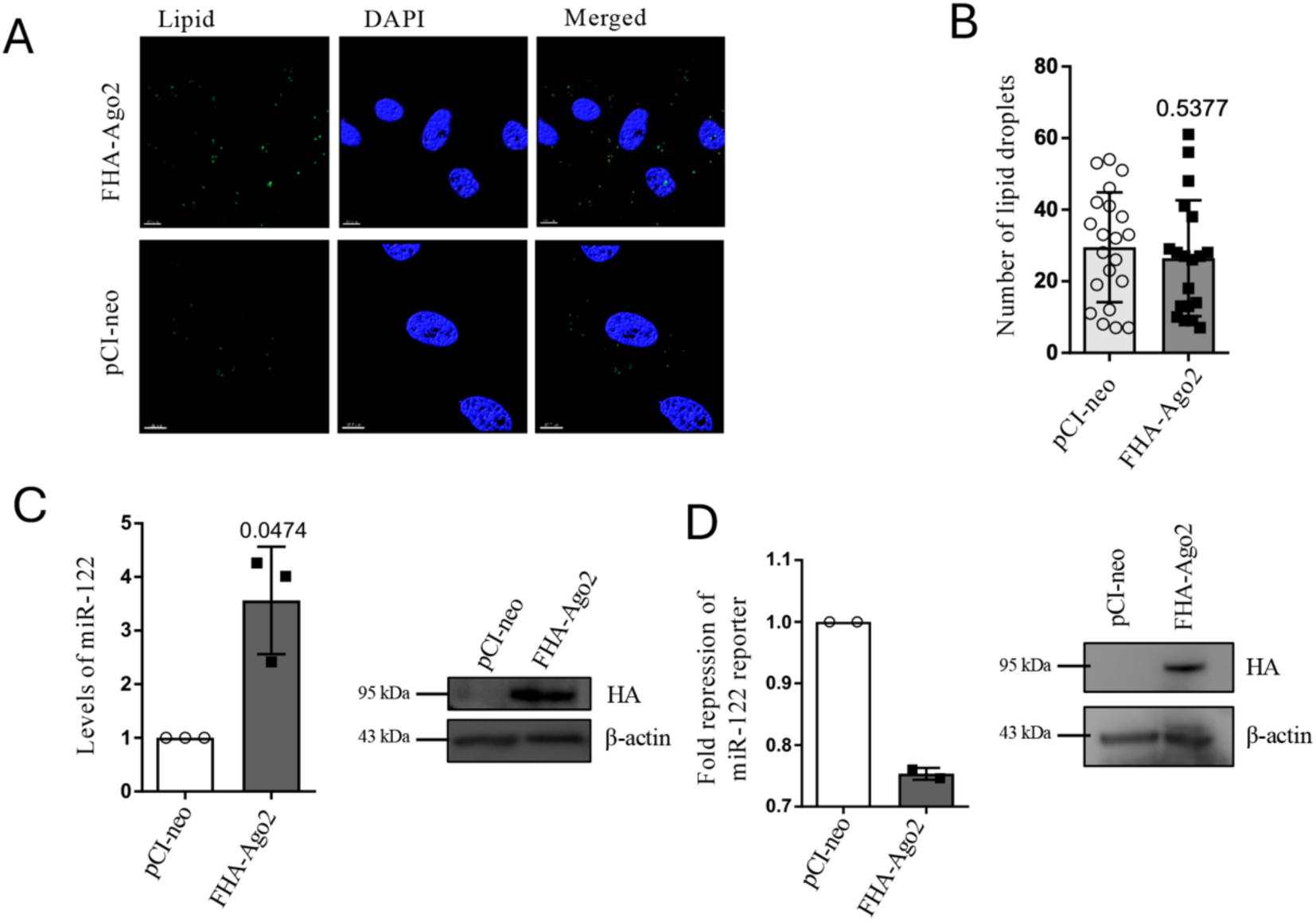
FHA-Ago2 affects miRNA content and activity. **A-B.** Effect of FHA-Ago2 expression on lipid droplets. LDs were visualized using the BODIPY 493/503 dye (green) in pCIneo- and FHA-Ago2-expressing cells co-transfected with 200nM miR-122 (A). Lipid droplet numbers were measured in 20 cells and plotted (B). Scale bar, 10 μm. **C-D.** Effect of FHA-Ago2 expression on miR-122 levels (C) and activity (D). PCIneo was used as a control. miRNA levels were estimated by qRT-PCR and normalized against U6. A Renilla luciferase reporter with miR-122 binding sites was used to measure miR-122 activity. FHA-Ago2 expression was confirmed by Western blot. All data are from three experimental replicates and are presented as SEM ± SD. P values are reported in the respective panels tested for statistical significance. P-values were calculated by a two-tailed paired t-test in most of the experiments unless mentioned otherwise. The fold change of miRNA was calculated by 2^−ΔCt^ method. The positions of the molecular weight markers are shown in the Western blot panel.

**Figure S5:**
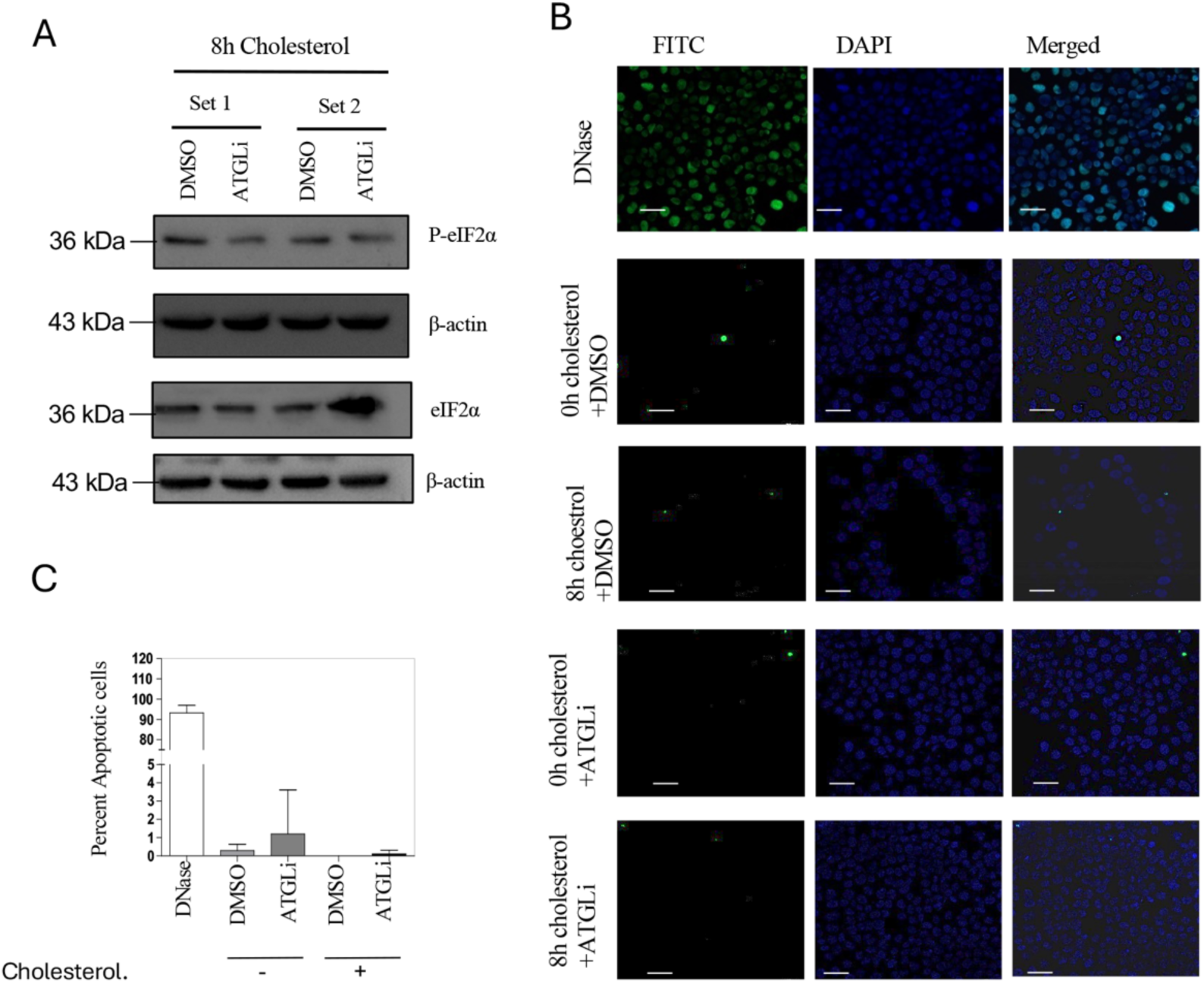
Enhanced lipid droplet number in cells with lipotoxicity reduces the stress response markers. A. The change in levels of P-eIF-2α in cells treated with ATGLi in the presence of 8h with MCD cholesterol concentrate. B-C. No change in apoptotic cell number with treatment of Huh7 for 8h with MCD-cholesterol concentrate. Apoptotic cells detected by TUNEL assay and respective quantification of percent TUNEL positive cells (B)/ The quantification against control DNAse-treated samples suggests a minimum number of apoptotic cells after 8h of MCD-Cholesterol treatment with ATGLi (10μM, 24h) treated Huh7 cells. All data are from three experimental replicates and are presented as SEM ± SD. The positions of the molecular weight markers are shown in the Western blot panel.

**S1 Table.**
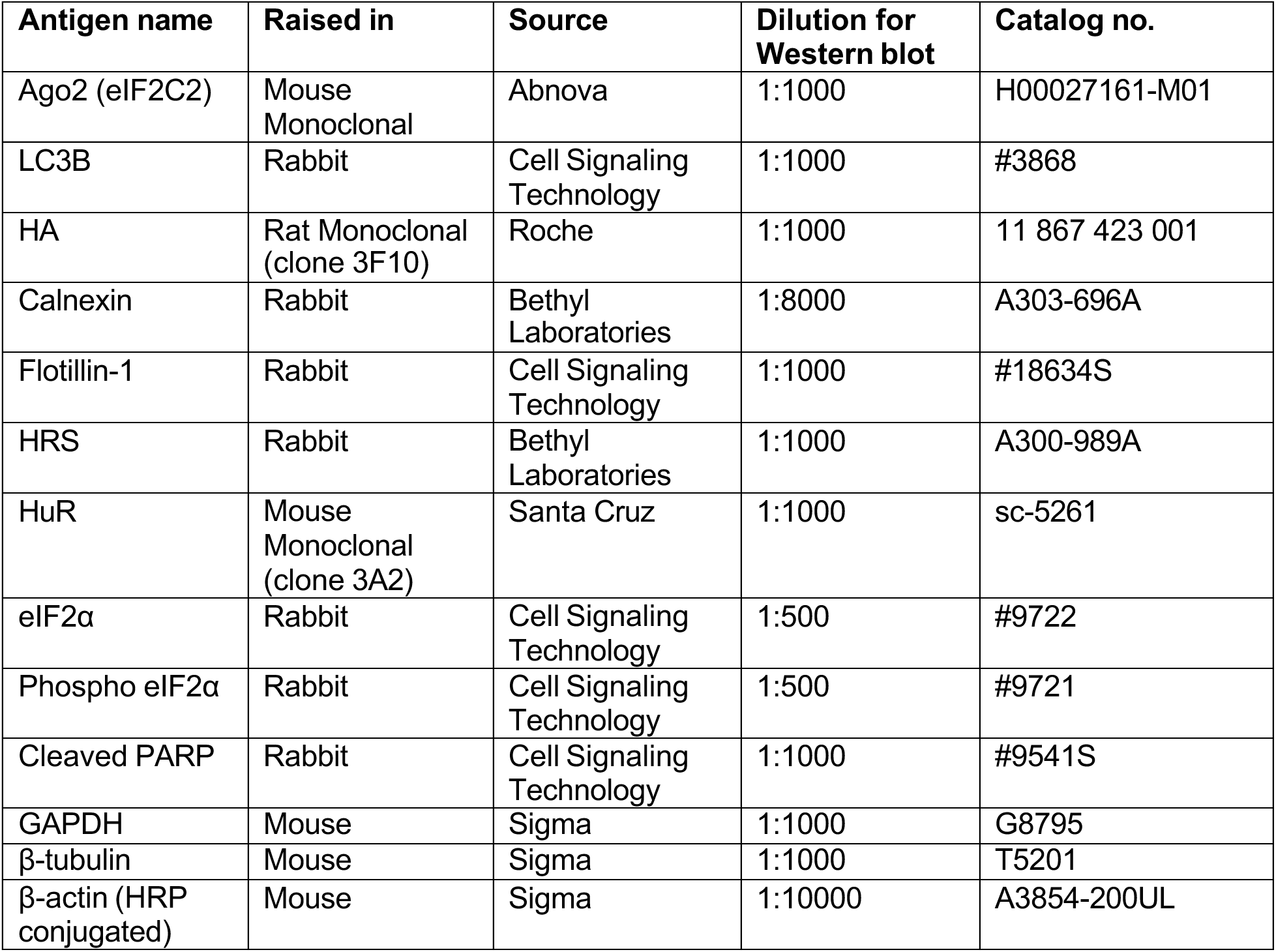
List of primary antibodies used.

| **Antigen name** | **Raised in** | **Source** | **Dilution for Western blot** | **Catalog no.** |
| --- | --- | --- | --- | --- |
| Ago2 (eIF2C2) | Mouse Monoclonal | Abnova | 1:1000 | H00027161-M01 |
| LC3B | Rabbit | Cell Signaling Technology | 1:1000 | #3868 |
| HA | Rat Monoclonal (clone 3F10) | Roche | 1:1000 | 11 867 423 001 |
| Calnexin | Rabbit | Bethyl Laboratories | 1:8000 | A303-696A |
| Flotillin-1 | Rabbit | Cell Signaling Technology | 1:1000 | #18634S |
| HRS | Rabbit | Bethyl Laboratories | 1:1000 | A300-989A |
| HuR | Mouse Monoclonal (clone 3A2) | Santa Cruz | 1:1000 | sc-5261 |
| eIF2α | Rabbit | Cell Signaling Technology | 1:500 | #9722 |
| Phospho eIF2α | Rabbit | Cell Signaling Technology | 1:500 | #9721 |
| Cleaved PARP | Rabbit | Cell Signaling Technology | 1:1000 | #9541S |
| GAPDH | Mouse | Sigma | 1:1000 | G8795 |
| β-tubulin | Mouse | Sigma | 1:1000 | T5201 |
| β-actin (HRP conjugated) | Mouse | Sigma | 1:10000 | A3854-200UL |

**S2 Table.** List of secondary antibodies used.

| **Secondary Antibody** | **Source** | **Dilution for Western blot** | **Catalog no.** |
| --- | --- | --- | --- |
| HRP Goat anti-mouse IgG | Life Technologies | 1:8000 | 626520 |
| HRP Goat anti-Rabbit IgG | Life Technologies | 1:8000 | 656120 |
| HRP Goat anti-Rat IgG | Life Technologies | 1:8000 | 629520 |

**S3 Table.**
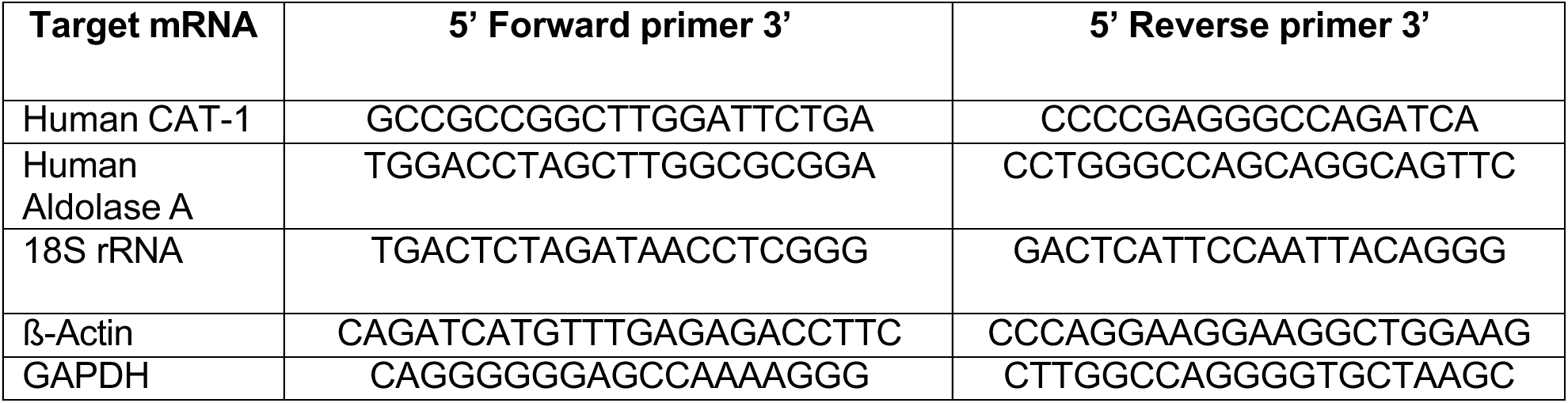
List of primers used for RT-qPCR detection of mRNAs.

**S4 Table.** List of primers used for RT-qPCR detection of miRNAs.

| Target miRNA | Assay ID |
| --- | --- |
| miR-122 | 002245 |
| miR-16 | 000391 |
| miR-21 | 000397 |
| let-7a | 000377 |
| U6 snRNA | 001973 |

**S5 Table.** List of Plasmids and siRNAs used.

| Name of Plasmid or siRNA | Source/ Reference | Plasmid Description |
| --- | --- | --- |
| FH-Ago2 | From Witold Filipowicz. Previously described in (1) | Plasmid expressing FLAG and HA tagged human Ago2 |
| pCI-neo | From Promega | pCI-neo expression vector |
| HA-HuR | From Witold Filipowicz. Previously described in (2) | Plasmid expressing HA tagged HuR, cloned in pCI-neo vector |
| GFP-PERILIN2 | From ORIGEN | Plasmid expressing GFP-tagged Perilin2 cloned in pCMV6-AC-GFP |
| siControl | Dharmacon | ON-TARGETplus Non-targeting Control pool |
| siPERILIPIN2 | Dharmacon | ON-TARGETplus SMARTpool- Human PERILIPIN2 |
| siHuR | Dharmacon | ON-TARGETplus SMARTpool- Human ELAV1 |

